# Long-term cellular and molecular signatures of pregnancy in the adult and ageing brain

**DOI:** 10.1101/2023.02.24.529879

**Authors:** P Duarte-Guterman, JE Richard, SE Lieblich, RS Eid, Y Lamers, LAM Galea

## Abstract

Pregnancy is marked by brain changes to volume, structure, connectivity, some of which are long-lasting. Few studies have examined possible mechanisms of these changes or the effects of multiple pregnancies. Here, we characterized various cellular and molecular signatures of parity (nulliparous, primiparous, biparous) in the hippocampus, an important area for cognitive and emotional regulation, and in plasma.

We investigated density of neural stems cells (Sox2) and microglia (Iba-1), and levels of the postsynaptic density protein (PSD-95), cell signalling pathways, hippocampal and peripheral inflammation and the tryptophan-kynurenine (TRP-KYN) pathway, at 1 week after weaning (7 months) and in middle-age (13 months). Parity increased PSD-95 levels in both age groups and prevented the age-related decrease in neural stem cell density observed in nulliparous rats. Biparity increased cell signalling phosphoproteins (pp706sk, S6RP) and number of microglia in the dentate gyrus, regardless of age. Parity resulted in transient changes to the TRP-KYN system and peripheral inflammation. Thus, parity has lasting effects on synaptic plasticity and alters the trajectory of hippocampal aging.

**Highlights:**

- Parity increased the postsynaptic protein PSD-95 in the hippocampus, regardless of age.
- Biparity increased microglial density and cell signalling in the hippocampus, regardless of age.
- Parity prevented the age-related decline in hippocampal neural stem cells.
- Parity transiently increased tryptophan-kynurenine pathway metabolites.
- Aging reduced plasma cytokine levels, an effect more prominent with nulliparity.

## 1. Introduction

During pregnancy, the entire body undergoes a massive physiological transformation. The placenta secretes large amounts of steroid and peptide hormones at levels not experienced outside of pregnancy, as progesterone is 20 times higher, and estradiol 200-300 times higher than during non-pregnant phases (Abbassi-Ghanavati et al., 2009; Levitz and Young, 1977). In addition, the immune, respiratory, metabolic, and cardiac systems adapt to allow for the growth of the fetus posing varied physiological challenges to the pregnant individual (Chauhan and Tadi, 2022; Napso et al., 2018; Pascual and Langaker, 2021). Indeed, pregnancy is a stress test for cardiovascular function (Facca et al., 2018) as cardiac output increases up to 80% (Chauhan and Tadi, 2022; Pascual and Langaker, 2022). The maternal brain also undergoes substantial remodeling during pregnancy, evident in the short- and long-term in both human and rodent studies (Eid et al., 2019; Hoekzema et al., 2017 and reviewed in Duarte-Guterman et al., 2019).

In primigravid individuals, i.e. individuals who are pregnant for the first time, total brain volume is reduced at parturition, but rebounds approximately 6 months postpartum (Oatridge et al., 2002). Multiple brain regions are remodelled during pregnancy with some areas rebounding (Carmona et al., 2019; Hu and Becker, 2003; Kim et al., 2018, 2010) but others, such as regions within the prefrontal cortex, showing enduring changes present years later (Hoekzema et al., 2017). Among the brain regions that are remodelled during pregnancy is the hippocampus, an important region for learning and memory and the regulation of stress and anxiety. The hippocampus is reduced in volume after giving birth and does not fully recover within 6 years after parturition (Hoekzema et al., 2017; Martínez-García et al., 2021a). An important consideration is that a reduction in volume need not be associated with impaired function, but instead could indicate fine tuning of the maternal brain to the demands of motherhood (Barha and Galea, 2017; Hoekzema et al., 2017; McCormack et al., 2023), and work in animal models indicates a facilitation of hippocampus-related cognition after the pups have weaned (Pawluski et al., 2006). Overall, motherhood-related changes to the brain vary depending on region, different stages of postpartum, years after birth and potentially amount of parity.

Most studies in humans concentrate on changes in the brain in the short-term, comparing pre-pregnancy to shortly after giving birth, or from early to late postpartum (2-6 months after giving birth). These studies show general trends of reductions in grey matter that occur during pregnancy with enhancements in grey matter from the early to later postpartum (Kim et al., 2018, 2010; Lisofsky et al., 2019; Luders et al., 2020, 2018; Martínez-García et al., 2021b; Rehbein et al., 2021). However, there are long-lasting signatures of previous pregnancy in the aging brain, as in middle and older aged women, previous parity is associated with less evident brain aging, including in the hippocampus (Lange et al., 2020; Orchard et al., 2020). Thus, findings suggest that there are long-lasting effects of parity in the brain, but few studies have examined the signatures at the cellular level that may act as possible mechanisms of these changes.

The underlying processes responsible for structural and connectivity changes during pregnancy and beyond are not completely understood and may include changes in the number of neurons and synapses. The hippocampus is unique as it shows a remarkable amount of plasticity with the continual production of new neurons (neurogenesis) throughout life in many mammalian species, including humans (Amrein, 2015; Kempermann et al., 2018; Zhou et al., 2022) as well as changes in dendritic architecture (spines/synapses, branching). In rats, adult hippocampal neurogenesis, spine density, dendritic branching and synaptic proteins change both in the short and long-term following pregnancy (Eid et al., 2019; Galea et al., 2018; Leuner et al., 2010; Pawluski et al., 2010; Pawluski and Galea, 2007; Workman et al., 2016). Collectively these results show that parity results in long-lasting changes to synaptic plasticity in multiple regions of the hippocampus that vary with age and amount of parity.

Pregnancy generates an anti-inflammatory milieu in the periphery to sustain the fetus and prevent fetal rejection (Robinson and Klein, 2012), although this depends on the trimester. The neuroimmune system also adapts during pregnancy and in the postpartum (Aagaard-Tillery et al., 2006; Haim et al., 2017; Luppi, 2003; Mor and Cardenas, 2010; Schumacher et al., 2014; Sherer et al., 2018), but few studies have investigated if these persist long term (Eid et al., 2019; Galea et al., 2018; reviewed in Barth and de Lange, 2020). Microglia, the resident immune cells in the brain, regulate many aspects of neuronal physiology including neurogenesis, synaptogenesis, and synaptic pruning (Haim et al., 2017; Hong et al., 2016; Sato, 2015; Wang et al., 2020). In the hippocampus, the number of microglia are reduced and levels of certain cytokines (interleukin 6 (IL-6) and IL-10) are increased during pregnancy and early postpartum (Eid et al., 2019; Haim et al., 2017; Posillico and Schwarz, 2016; Sherer et al., 2018, 2017) and some of the changes in microglia and peripheral cytokines can persist in the long term (Eid et al., 2019).

Pregnancy-induced changes to neuroimmune signalling may therefore cause long-lasting changes to hippocampal function, although little is known about the how these transient changes in immune signalling affect brain health and processing beyond pregnancy and postpartum. Furthermore, little is known about the short- and long-term neuroimmune consequences of increasing amount of parity.

Pregnancy also alters the tryptophan (TRP)-kynurenine (KYN) pathway (van de Kamp and Smolen, 1995), which has a direct relationship with the inflammatory system, as proinflammatory cytokines, such as IFNγ and IL-1β, increase enzymatic conversion of tryptophan into its downstream metabolite, KYN (Maes et al., 2011). In addition, the KYN pathway is altered with age, and alterations in its activity are associated with a variety of age-associated neurological diseases (Castro-Portuguez and Sutphin, 2020).

Branches of the TRP-KYN pathway can be divided into either “neurotoxic” or “neuroprotective” in which KYN has neurotoxic properties whereas kynurenic acid (KA) is thought to be neuroprotective (Wichers et al., 2005). Pregnancy increases TRP utilization to support fetal growth and development, provide increased levels of serotonin for signaling pathways, KA for neuronal protection and other kynurenines to suppress fetal rejection (Badawy, 2015). The TRP-KYN pathway also regulates hippocampal neurogenesis and inflammation, (Hasegawa et al., 2010; Jung et al., 2020; Zysset-Burri et al., 2013).

Several key enzymes of the TRP-KYN pathway require vitamin B6, and deficiency develops gradually during the course of pregnancy (Reinken and Dapunt, 1978; van de Kamp and Smolen, 1995). To our knowledge no study has investigated the longer-term effects of pregnancy and motherhood on the TRP-KYN pathway.

Our objective was to perform a wide-ranging study of the cellular and molecular signatures of parity (pregnancy and motherhood experience) in the hippocampus after weaning and in middle-age. We investigated the effect of nulliparity (no pregnancy or motherhood experience), primiparity (pregnant and mothers to one litter), and biparity pregnant twice and mothers to two litters) in the dorsal and ventral hippocampus. We measured levels of the synaptic protein post-synapse density (PSD-95), ribosomal protein S6 kinase (pp70S6K) and S6 ribosomal protein (S6RP), neuroinflammation markers (cytokines, chemokines), metabolites in the TRP-KYN pathway, and cell signalling proteins of the ERK, and Akt pathways involved in a range of functions including synaptic protein transport and immune signalling (Yoshii and Constantine-Paton, 2014). We also examined neural stem cells (Sox2+ cells) and microglia (Iba-1). We expected that parity would increase neuroinflammation and reduce neuroplasticity in the short term, and that these factors would normalize or reverse (increased synaptic proteins) long-term compared to nulliparous females, which would be dependent on the amount of parity.

## 2. Material and methods

### 2.1. Animals

Adult male and female Sprague–Dawley rats (2 months old) were obtained from Charles River (Quebec, Canada). Rats were given *ad libitum* access to food and water and double-housed, except during gestation and in the postpartum until weaning in which they were single-housed (nulliparous females were single-housed for an equivalent period). Nulliparous rats were housed in a separate colony room during breeding and until weaning was completed to prevent nulliparous females from being exposed to males and to any offspring. All protocols were approved by the Institutional Animal Care Committee at the University of British Columbia and conformed to the guidelines set out by the Canadian Counsel on Animal Care.

### 2.2. Breeding and maternal observations

For breeding, two females and one male were paired overnight. Females were vaginally lavaged each morning, and samples were assessed for the presence of sperm. If sperm was present, the female was considered pregnant, weighed, single-housed into a clean cage, and monitored weekly throughout gestation. One day after birth (postpartum day 1), litters were culled to 4 males and 4 females. One female from the young adult biparous group did not have enough pups, therefore we cross-fostered with pups from other females born on the same day. All animals were age-matched such that primiparous rats were the same age as biparous rats during their first and second pregnancy, respectively. Biparous rats were bred at 3 months and 5 months of age and primiparous females were bred at 5 months of age. Maternal behavior was assessed. Each observation per animal lasted for 20 min per day (10 min per morning and afternoon session). The following maternal care behaviors were scored: nursing, which included arched-back nursing (arched-back, actively nursing pups), blanket nursing (in nursing position but resting on-top of pups), and passive nursing (lying on the side and nursing pups); licking (licking pups while not nursing any other pups), self-grooming (not engaging with pups, and self-grooming), and time spent off-nest (off-nest and not engaging with pups).

Timeline is illustrated in **Fig. 1**.

**Figure 1.**
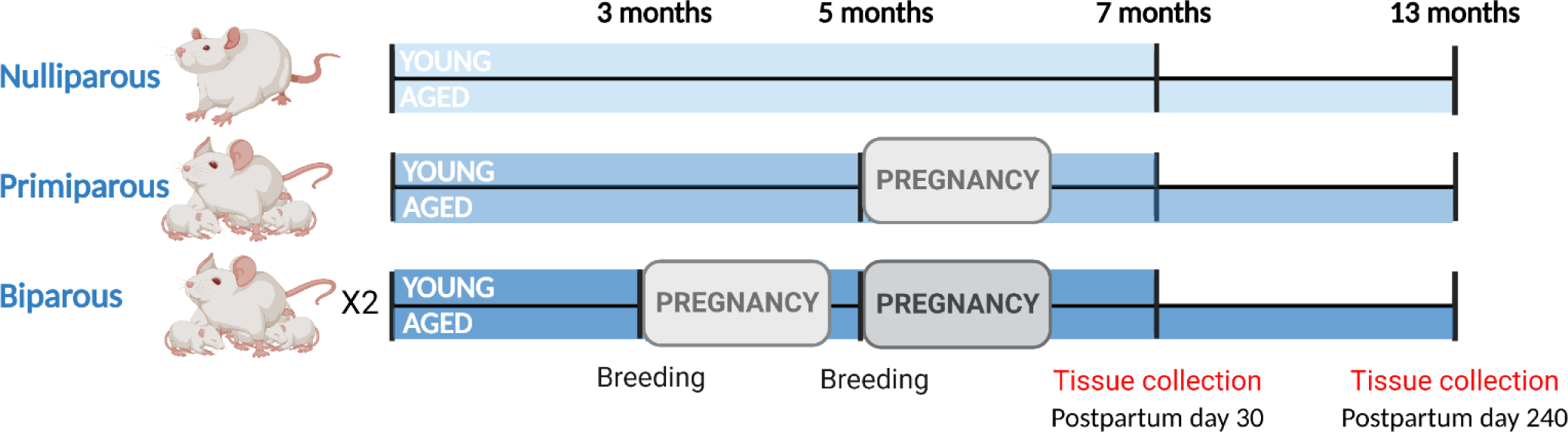
Graphical representation of experimental timeline, including pregnancies and tissue/sample collection. Image created using BioRender.com.

### 2.3. Blood and Tissue Collection

To monitor estrous cycles, vaginal lavages were conducted for 5-7 days before tissue collection (starting postpartum day 23) to determine whether cycling had re-started after pregnancy in the young adult animals or had stopped in the older animals (cycling data is reported in **Fig. 2C**). Lavages were also conducted on day of euthanasia. On postpartum day 30 (adult) and 240 (middle-aged) all rats received a lethal overdose of sodium pentobarbital between 10 am-12 pm, to limit any circadian influence. Blood was collected via cardiac puncture into cold EDTA-coated tubes and centrifuged (within 2 h from time of dissection) for 10 min at 4°C and plasma was stored at -80°C. Brains were quickly extracted, and hippocampus was rapidly dissected from the left hemisphere over ice, flash frozen on dry ice, and stored at -80°C (n=10/group). Cytokine levels in the hippocampus were analyzed using electrochemiluminescence immunoassay kits. The right hemisphere (n=3-6/group) was fixed for 24 h in 4% paraformaldehyde in PBS (4°C) and then transferred to a 30% sucrose/PBS solution for cryoprotection. Serial coronal sections (40 µm) were cut with a freezing microtome (SM2000R; Leica, Richmond Hill, ON) across the extent of the hippocampus (collected in 10 series) and stored in an antifreeze solution (containing 0.1M PBS, 30% ethylene glycol, and 20% glycerol) and stored at -20°C for immunohistochemistry.

**Figure 2.**
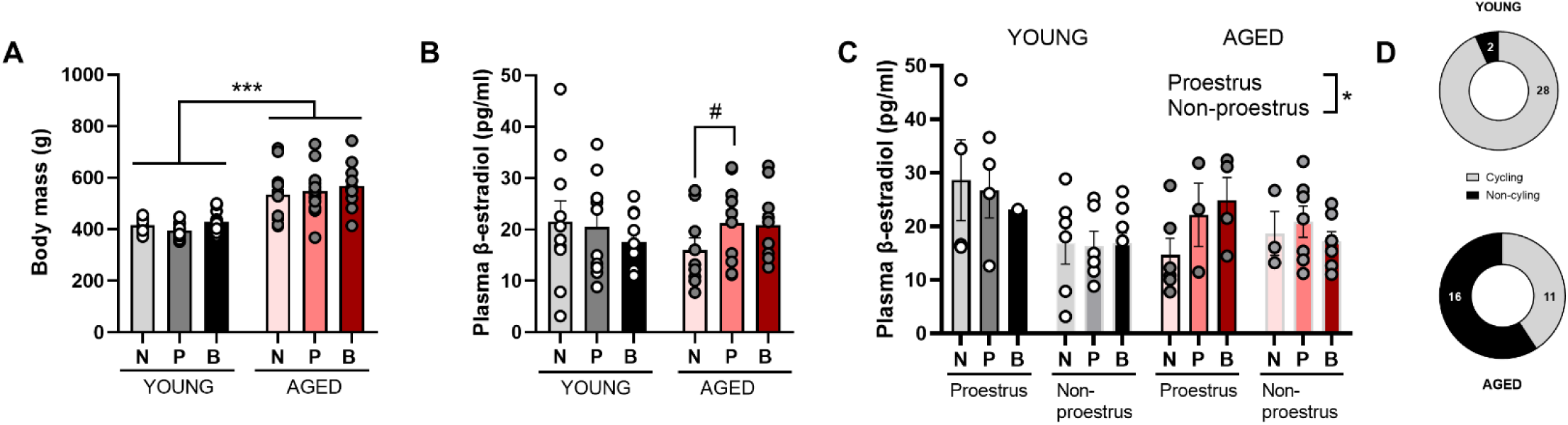
Body mass increases in middle-aged regardless of parity and serum β-estradiol is higher in proestrus than non-proestrus cycle phases. Female rats displayed increased body mass in all parity groups (A). There is a trend towards increased levels of plasma estradiol in aged primiparous, compared to aged nulliparous, females (B). Plasma β-estradiol levels are higher in proestrus than non-proestrus (C). Proportion of cycle and non-cycling females in each age group are presented in (D). Presence of cycling in 3 females in the aged group could not be determined. Data are expressed as mean ±SEM. Body mass data: n = 9-10 per group; β-estradiol data: n=5-21 (merged), and n=1-9 (by parity) per group. * p < 0.05, ** p < 0.01, *** p < 0.001, # p = 0.060.

### 2.4. Plasma hormone and TRP-KYN metabolite quantification

Estradiol levels were determined in plasma (in duplicate) using a radioimmunoassay kit (ultrasensitive estradiol RIA, Beckman Coulter, DSL4800; assay sensitivity is 2.2pg/mL, average intra- and inter-assay coefficients of variation were <20%).

We quantified KYN, KA, xanthurenic acid (XA), anthranilic acid (AA), 3-hydroxykynurenine (3-HK), 3-hydroxyanthranilic acid (3-HAA), riboflavin (vitamin B2) and its cofactors, flavin adenine dinucleotide (FAD) and flavin mononucleotide (FMN) from plasma, using isotope dilution liquid chromatography coupled with tandem mass spectrometry (ABSciex API4000; AB SCIEX, Framingham, MA, U.S.A.) based on a modified method by (Midttun et al., 2005). Neuroprotective-to-neurotoxic z score ratios were calculated to account for the balance between neuroprotective metabolites or neurotoxic metabolites/cofactors: zKA/zHA and zKA/zHK. A similar approach was used in (Qiu et al., 2021).

### 2.5. Hippocampal PSD-95, cytokines, and cell signaling levels

Electrochemiluminescence immunoassay kits (Meso-Scale Discovery; Rockville, MD) were used to measure PSD-95 (Cat# K150QND), ERK-1/2, JNK, p38 (MAP Kinase Whole Cell Lysate Kit; Cat # K15101D), Akt, GSK-3β, p70S6K, S6RP (Akt Signaling Panel II Whole Cell Lysate Kit; Cat# K15177D) and cytokines (IFN-γ, IL-1β, IL-4, IL-5, IL-6, IL-10, IL-13, CXCL-1, and TNF-α; V-PLEX Proinflammatory Panel 2 Rat Kit; Cat# K15059D) in homogenized whole hippocampal samples. Levels of MEK 1 and 2, and STAT3 (Cat# K15116) were also measured, although levels were undetectable. Hippocampi were homogenized using a bead disruptor (Omni, VWR) in 200 µl of cold Tris lysis buffer (150 mM NaCl, 20 mM Tris, pH 7.5, 1 mM EDTA, 1 mM EGTA, 1% Triton X) containing a cocktail of protease and phosphatase inhibitors. Homogenates were centrifuged at 2100g for 10 min at 4°C and supernatants were aliquoted and stored at -80°C. Protein concentrations were determined using a BCA protein assay kit (ThermoFisher). Homogenates were diluted with Tris lysis buffer 1:10 for PSD-95 and Akt kit plates and 1:100 for MAP kinase kit. Samples were run in duplicate following the manufacturer’s protocols except that plates were incubated overnight at 4°C (instead of 3 h). Plates were read with a MESO QuickPlex SQ 120 (Meso Scale Discovery) and data were analyzed using the Discovery Workbench 4.0 software (Meso Scale Discovery). Cytokine levels were obtained using a standard curve and expressed as pg/mg protein. For PSD-95 and cell signaling assays, background signal was subtracted from raw signal and normalized to protein levels and results are expressed as signal/mg/ml protein.

### 2.6. Iba1 immunohistochemistry

A series of free-floating tissue was rinsed 3x with 0.1 M PBS for 10 min each. Tissue was then incubated in 3% hydrogen peroxide for 30 mins. Sections were added to the blocking solution (10% normal goat serum (NGS) and 0.4% Triton X 100 in PBS) for 1 h. Tissue was immediately incubated in a primary antibody solution (1:1000 rabbit anti-Iba1 (Wako Chemicals USA) in PBS containing 5% NGS, 0.4% Triton X) for ∼20 h at 4°C. Following PBS rinses, tissue was incubated for 1.5 h in a biotinylated goat anti-rabbit antibody (1:500; Vector laboratories) in PBS containing 2.5% NGS, 0.4% Triton-X. Sections were incubated in avidin-biotin horseradish peroxidase complex kit (1:500, Vector Laboratories) in PBS containing 0.4% Triton-X for 1 h. To visualize the Iba1 positive cells, tissue was incubated in 3,3’ - diaminobenzidine Peroxidase Substrate Kit (Vector Laboratories) using 0.175M sodium acetate as the buffer for ∼4 min. Tissue was rinsed 3x quickly, then another 3x 10 min each to stop the reaction. Finally, tissue was mounted onto Superfrost/Plus slides (Fisher scientific), allowed to dry a few days, then dehydrated in ethanol, cleared with Xylene, and cover slipped immediately with Permount (Fisher Scientific). The primary antibody was omitted as a control and tissue did not contain any immunoreactivity.

### 2.7. Sox2 immunohistochemistry

A series of free-floating tissue was rinsed 3x with 0.1M tris buffered saline (TBS; pH 7.4) for 10 min each, on a rotator at room temperature (RT), between each of the following steps unless otherwise stated. Sections were incubated in 3% hydrogen peroxide for 30 min RT. Tissue was blocked in a TBS solution containing 3% normal horse serum (NHS) and 0.3% Triton X 100 for 30 min RT. Tissue was immediately incubated in a primary antibody solution (1:1000 mouse anti-Sox2 in TBS containing 3% NHS, 0.4% Triton X) ∼40 h at 4°C. Following TBS rinses, tissue was incubated for 2 h with ImmPRESS HRP Horse anti-Mouse IgG, (1:2, Rat adsorbed, Vector laboratories) diluted in TBS. To visualize Sox2 positive cells, tissue was incubated in 3,3’ -diaminobenzidine Peroxidase Substrate Kit plus Nickel per the Vector lab kit instructions for ∼7.5 min. Tissue was rinsed 3x quickly, then another 3x 10 min each to stop the reaction. Tissue sections were mounted onto Superfrost/Plus slides (Fisher scientific), allowed to dry, then dehydrated in ethanol, cleared with Xylene, and coverslipped immediately with Permount (Fisher Scientific). The primary antibody was omitted as a control and tissue did not contain any immunoreactivity.

### 2.8. Microscopy analysis

A researcher blind to experimental conditions counted Iba1+ cells in one hemisphere for the entire rostrocaudal (dorsal and ventral) extent of the granule cell layer (GCL) at 400x magnification using a standard light microscope. GCL volumes were quantified from digitized images using Cavelier’s principle, multiplying the sum of the area of each section by the section thickness (30 µm; (Gundersen, 1988). Counts were divided by GCL volume per section to obtain cell densities (counts/mm^3^). Additionally, we examined Iba-1+ cell morphology as a proxy for functional state, following previously described methods (Eid et al., 2019). Briefly, an experimenter blinded to condition categorized Iba1+ cells into ramified (highly branched with long processes), stout few, shorter processed), or ameboid microglia (no processes). For the CA1 and CA3 regions of the hippocampus, photomicrographs of three dorsal and three ventral sections (6 sections total per animal) were taken using a 10X objective and the same exposure and gain settings. For the CA1 and CA3, Iba-1+ cell counts were divided by the area of the region of interest (counts/mm^2^). Representative image of Iba1+ cells in the hippocampus demonstrated in Fig 4D-E at 4X.

Sox2+ cells were counted in the SGZ and GCL of the dentate gyrus in five dorsal and five ventral sections (10 sections total per animal) using segmentation and object classification analysis in the density counting workflow in Ilastik (Berg et al., 2019). Photomicrographs were taken at 40X magnification. Dorsal and ventral regions were trained separately due to differences in background. Five (dorsal) and six (ventral) images were used for training to define objects (Sox2+ cells) and background. Images for training were chosen from the two ages (7 and 13 months) and all parity groups. Images were exported to ImageJ for tracing the GCL and SGZ regions on each section and density of Sox2+ cells were obtained using the integrative density function. Representative images of Sox2+ staining in the hippocampus are demonstrated in Fig 4A-B.

### 2.9. Statistical analysis

All the data are presented as mean ± standard error of the mean. Data were analyzed using TIBCO Statistica software (v. 9, StatSoft, Inc., Tulsa, OK, USA) using ANOVA using parity and age as between subject factors and plotted using GraphPad Prism (GraphPad Software, Inc., San Diego, CA). Simple linear regression was used to analyze influence of sex litter ratio on cytokine levels. Effect sizes (partial eta squared (Ƞ_p_^2^) and Cohen’s *d*) are reported for significant effects. Post hoc comparisons used Newman-Keul’s. *A priori* we hypothesized that there would be differences between parity groups and any *a priori* comparisons were adjusted using Bonferroni correction. For cytokines and TRP-KYN metabolites principal component analyses (PCA) were conducted to determine networks (components) that could explain metabolite variances. Principal component scores for individual sample data for the first three components were generated using R Statistical Software (v4.1.2; R Core Team 2021) and analyzed using two-way ANOVA models to derive information regarding the amount of variance accounted for by metabolite and cytokine profiles. Frequency of cycling between the age groups was compared using Pearson’s chi-square (χ^2^) test. P-values lower than 0.05 were considered statistically significant.

## 3. Results

### 3.1 Increased age is associated with increased body mass and reduced cycling

The biparous dam spent significantly less time licking pups (F_(1,62)_ = 4.633, *p* = 0.035) and self-grooming (F_(1,62)_ = 4.275, *p* = 0.043) in the second litter compared to the first litter (data not shown). There were no other significant differences in the other maternal behaviors (*p*’s > 0.461).

As expected, middle-aged females had higher body mass than young adult females (main effect of age, F_(1, 52)_ = 50.022, *p =* 0.000, Ƞ_p_ = 0.490), regardless of parity (nulliparous vs nulliparous: *p =* 0.002, Cohen’s *d* = 1.574; primiparous vs primiparous: *p =* 0.0003, Cohen’s *d* = 2.168; biparous vs biparous: *p* < 0.0009, Cohen’s *d* = 1.998836). There was no significant main effect of parity (*p =* 0.527) and no significant interaction between parity and age regarding body mass (*p =* 0.759; **Fig. 2A**).

Next, we examined estradiol levels. Estradiol did not significantly vary depending on age or parity amount (all *p*’s > 0.268; **Fig. 2B**). However, there was a trend towards an increase in estradiol levels in nulliparous, compared to primiparous, aged females ( *p* = 0.060, Cohen’s *d* = 0.692). One outlier (Grubbs test) was removed from the primiparous aged group; this value was double the other females’ values.

There was a significant effect of cycle phase on estradiol levels (main effect cycle phase: F_(1,47)_ = 4.636, *p* = 0.036, Ƞ_p_^2^ = 0.090; **Fig. 2C**). More specifically, females in proestrus had higher levels of estradiol than females in other cycle phases ( *p* = 0.014, Cohen’s *d* = 0.421). There were no other significant main or interaction effects (*p*’s > 0.359). As expected, middle-aged rats were more likely to be non-cycling than younger adult rats (χ^2^  = 30.523, *p* = 0.00002; **Fig. 2D**). Parity did not affect likelihood of cycling (*p* > 0.541).

### 3.1 Parity increases synaptic plasticity and cell signaling phosphoproteins in the hippocampus, regardless of age

We investigated the effects of parity on cell signaling and synaptic proteins in the hippocampus. Parity significantly increased PSD-95, regardless of age (main effect parity: F_(2, 55)_ = 4.612, *p* = 0.014, Ƞ_p_ = 0.143; **Fig. 3A**), with both primiparous and biparous animals having more PSD-95 expression relative to nulliparous animals (p’s < 0.022, Cohen’s *d ≥* 0.813). There were no other significant main or interaction effects (*p* > 0.349).

**Figure 3.**
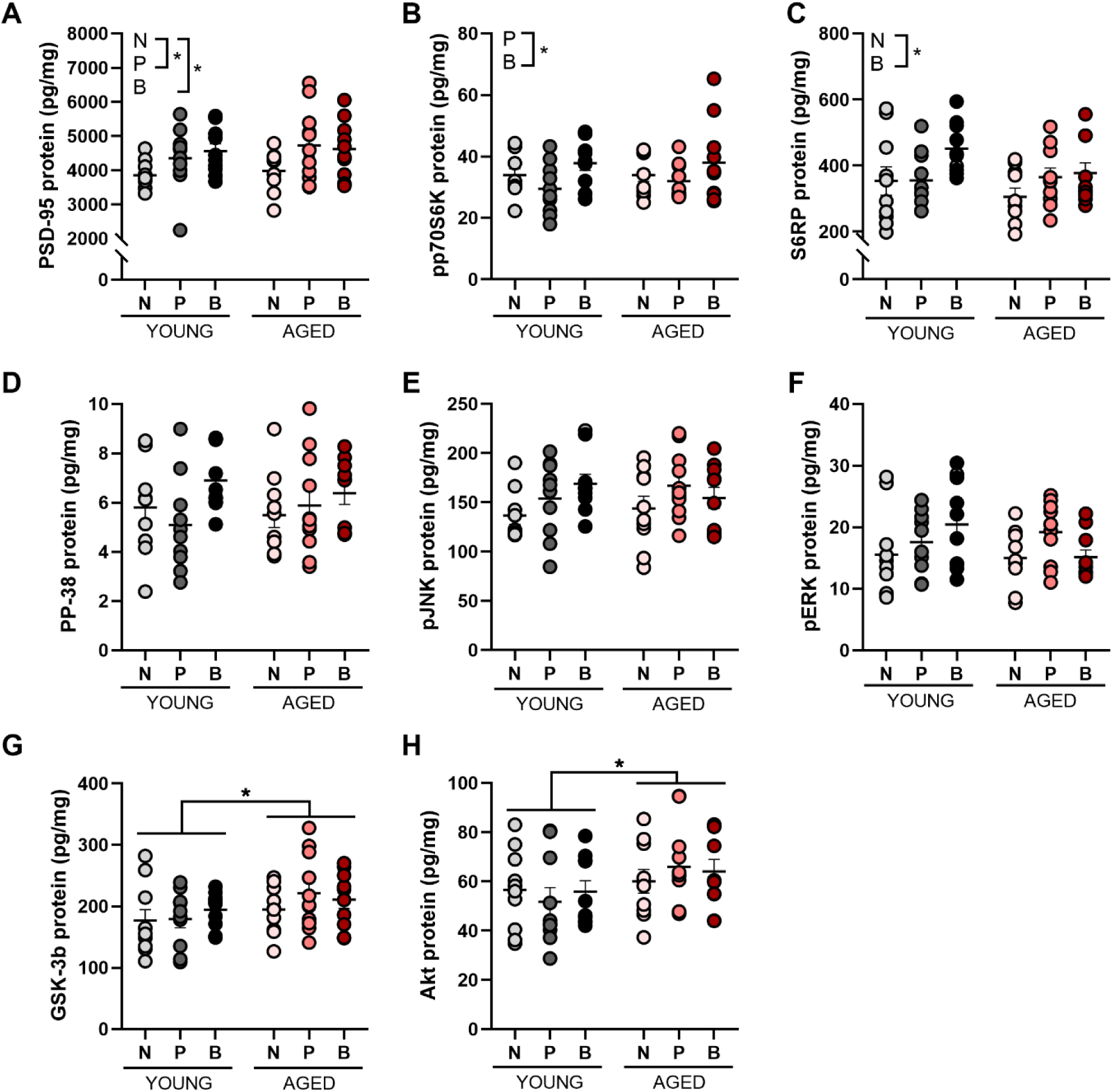
Parity increased synaptic plasticity in the hippocampus. Parity increased the synaptic protein PSD-95 (A), enzyme pp70S6K (B), and cell signalling phosphooprotein S6RP (C), regardless of age. Age significantly increased the levels of Akt (H) and GSK-3b (G). Parity did not significantly affect protein levels of PP-38 (D), pJNK (E), pERK (F), GSK-3b (G) and Akt (H). Data are expressed as mean ±SEM. n = 9-11 per group. * p < 0.05.

Biparity increased expression of the phosphoprotein ribosomal protein S6 kinase (pp70S6K) relative to the primiparous group (*p* = 0.020, Cohen’s *d* = 0.812) but not the nulliparous group (*p* = 0.122; main effect of parity: F_(2, 55)_ = 3.952, *p* = 0.025, Ƞ_p_ = 0.126; **Fig. 3B**). Biparity increased expression of the phosphoprotein S6RP relative to nulliparity (*p* = 0.019, Cohen’s *d* = 1.063) but not primiparity (*p =* 0.342; main effect of parity: F_(2, 55)_ = 2.959, *p* = 0.060, Ƞ_p_ = 0.097; **Fig. 3C**). There were no other main effects or interactions for these phosphoproteins (all *p*’s > 0.180).

Aging increased the expression of the phosphoproteins Akt and GSK-3b (main effect of age: Akt F_(1, 53)_ = 4.276, *p* = 0.044, Ƞ_p_^2^ = 0.075, and GSK-3b F_(1, 53)_ = 4.658, *p* = 0.035, Ƞ_p_^2^ = 0.081; **Fig. 3G** and **H**), with higher levels of these phosphoproteins in middle age, than in young adult females (Akt: *p* = 0.040, Cohen’s *d* = 54.798 and GSK-3b: *p* = 0.038, Cohen’s *d* = 0.552), but there were no other significant main or interaction effects (*p* > 0.431). There were no other main effects or interactions for any of the other phosphoproteins examined (all *p*’s > 0.137; **Fig. 3D-F**).

### 3.1 Aging reduced density of neural stem cells in nulliparous rats only and biparous rats had greater density of microglia in the dentate gyrus

Next, we investigated if parity alters the density of SOX2+ neural stem cells within the dentate gyrus (**Fig. 4A-B**). Although there was a main effect of age to reduce SOX2+ cell density (main effect age: F_(1, 24)_ = 5.614, *p* = 0.026, Ƞ_p_ = 0.078;), a priori comparisons indicated that SOX2 cell density was significantly reduced with age in nulliparous(*p* = 0.013, Cohen’s *d* = 0.978), but not in primiparous (*p* = 0.387) or biparous (*p* = 0.484) rats (**Fig. 4C**). There were no other significant effects (*p’s >* 0.379).

**Figure 4.**
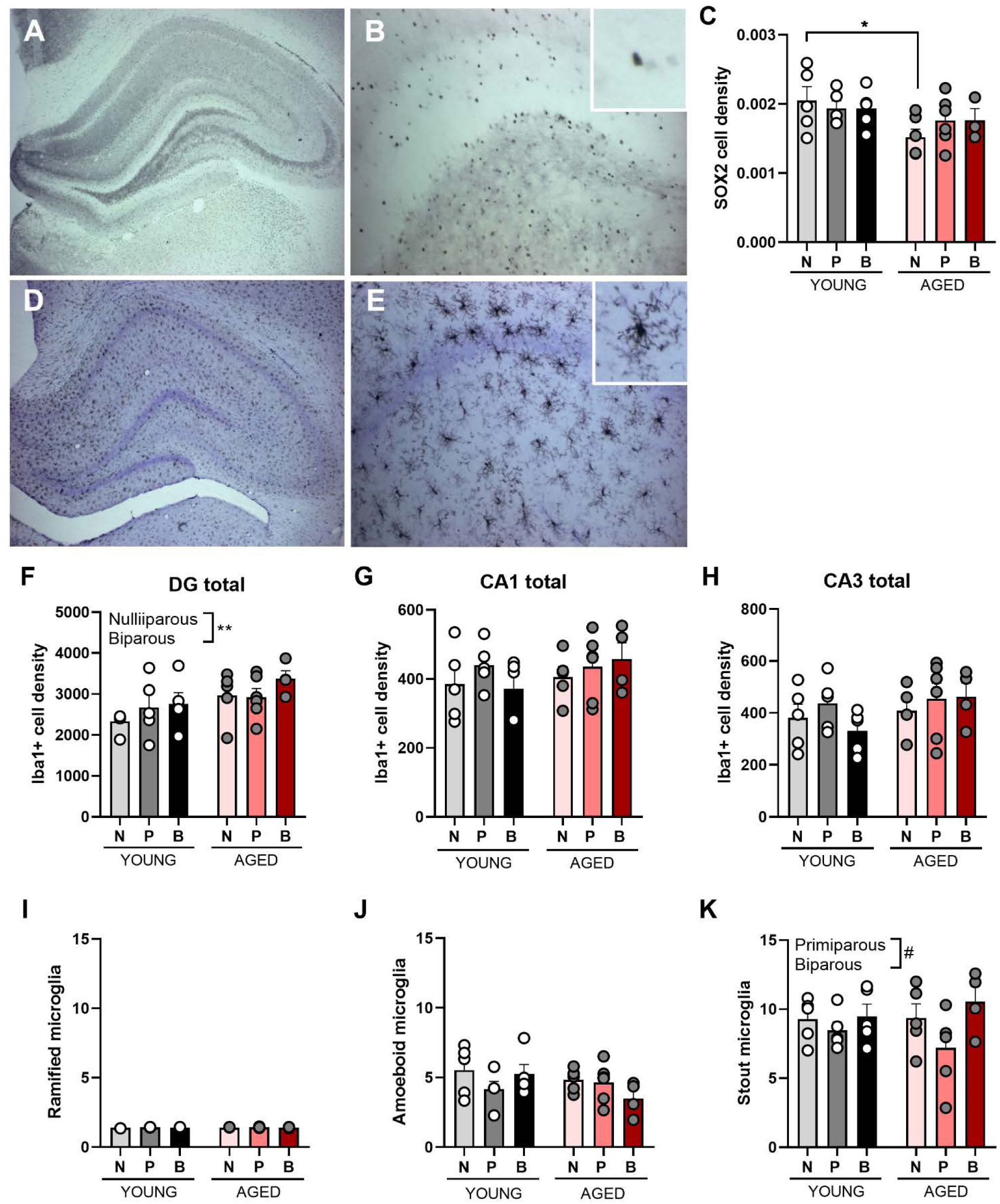
Increasing age decreases neural stem cells in nulliparous rats only, and increases microglia density in the hippocampus regardless of parity. Representative images of SOX2 positive cells at 4X (A) and 20X (B). SOX2 cell density is reduced with age in nulliparous, but not parous females (C). Representative image of Iba1+ positive cells at 4X (D) and 20X (E). Iba1+ cell density is significantly higher in biparous females, compared to nulliparous females, in the dentate gyrus (DG; F). No differences in the Iba1+ cell density was found in the CA1 (G) or CA3 (H) regions. Proportion of ramified (I), amoeboid (J) and stout (K) microglia (Iba1+ cells) in the GCL. Data are expressed as mean ±SEM. n = 4-6 per group for Iba1+ and 3-6 per group for SOX2. * p < 0.05, # = 0.071. N = nulliparous, P = primiparous, B = biparous.

Because immune processes impact hippocampal plasticity, we next investigated potential effects of age and parity on microglial number (representative image displayed in **Fig. 4D-E**). Middle-aged rats had more microglia in the dentate gyrus than younger adult rats (main effect of age: F_(1, 24)_ = 6.129, p = 0.021, Ƞp2 = 0.203). Biparous rats had a greater density of Iba1+ cells in the dentate gyrus compared to nulliparous (*p* = 0.001) rats (a priori comparisons; **Fig. 4F**), but not primiparous rats (*p* = 0.166), regardless of age. There were no other significant effects in the dentate gyrus (all *p’s* > 0.263) or in the CA3 or CA1 regions (all *p’s* > 0.16 see **Fig. 4G and H**).

As for Iba1+ cell morphology, there was a trend towards a reduction in stout Iba1+ cells in primiparous females, compared to biparous females (*p* = 0.071, Cohen’s *d* = 1.042; main effect of parity: F_(2, 24)_ =2.877, *p* = 0.076, Ƞ_p_^2^ = 0.193; **Fig. 4K**). There were no other significant effects (*p* > 0.510). There were no significant effects of parity on ramified or amoeboid Iba1+ cells (*p* > 0.112; **Fig. 4I-J**).

### 3.1 Minimal effects of ageing and parity on hippocampal cytokines, and plasma cytokine levels are mainly affected by age

There was a trend towards a reduction in hippocampal CXCL-1 levels in aged compared to young biparous females (*p* = 0.072, Cohen’s *d* = 0.855; interaction between age and parity: F_(2, 52)_ = 3.368, *p* = 0.083, Ƞ_p_ = 0.090; **Fig. 5I**). There was also a trend towards a significant main effect of parity for hippocampal IL-6 (IL-6: F_(2, 53)_ = 2.463, *p* = 0.095, Ƞ_p_ = 0.085; **Fig. 5D**). There were no significant effects for hippocampal IL-1β (*p* > 0.113, **Fig. 5A**), IL-4 (*p* > 0.366, **Fig. 5B**), IL-5 (*p* > 0.229, **Fig. 5C**), IL-10 (*p* > 0.299, **Fig. 5E**), IL-13 (*p* > 0.111, **Fig. 5 F**), IFN-γ (*p* > 0.132, **Fig. 5G**) or TNFα (*p* > 0.221, **Fig. 5H**). Cytokine levels were not influenced by sex litter ratio (all *p’s >* 0.677; *Supplementary Fig. 1*). We also analyzed the hippocampal cytokines using PCA. Component loading are displayed in Table 1. Results from the PCA analysis yielded two principal components, which together explained 80.98% of the total variance. All 9 cytokines examined loaded positively onto principal component analysis (PCA) 1 which explained 68.74% of the variance (**Table 1**). No significant effects between parity and age groups were demonstrated in PCA1 (*p’s* > 0.141; **Fig. 5J**).

**Figure 5.**
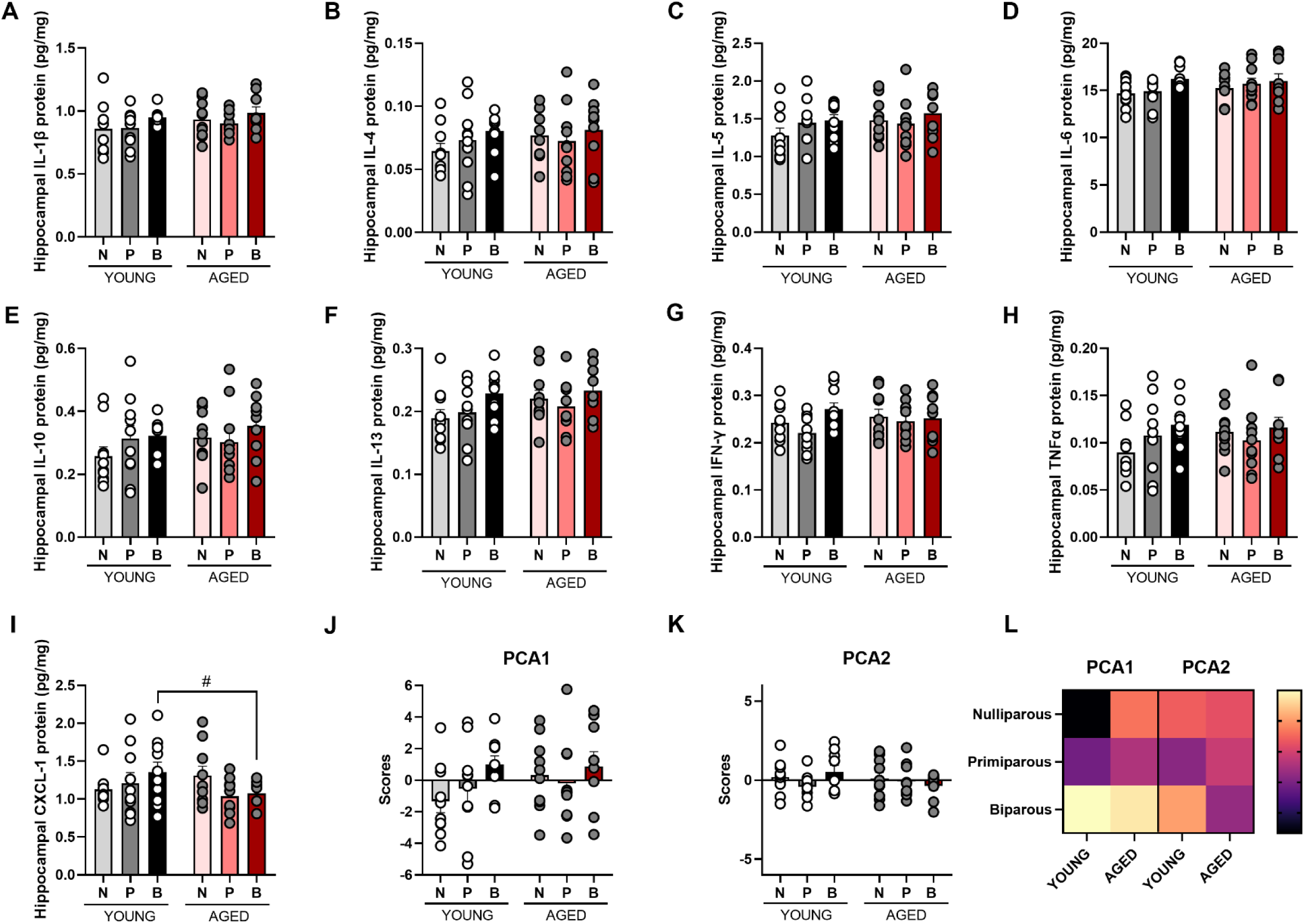
Levels of hippocampal cytokines in young adult and aged females of different parity. Protein levels of cytokines interleukin (IL)-1α (A), IL-4 (B), IL-5 (C), IL-6 (D), IL-10 (E), IL-13 (F), interferon (IFN)-γ (G), tumor necrosis factor (TNF)-α (H) and chemokine ligand (CXCL)-1 (I) measured in the hippocampus. # *p* = 0.072. Loadings of scores for each subject onto PCA 1 (J) and 2 (K). Heat map displaying average loading of each parity group on each compartment (L). No significant differences between parity or age groups were found. Expression data are presented in pg/mg and all data is expressed as mean ±SEM. n = 9-10 per group N – nulliparous, P = primiparous, B = biparous.

**Table 1.** Component loadings of the hippocampal cytokine variance of components identified using principal component analysis (PCA).

| **PCA1** | **Cytokine (hippocampus)** | **Correlation** | **p-value** |
| --- | --- | --- | --- |
|  | IL-13 | 0.9541556 | **1.51e-31** |
|  | TNFalpha | 0.9344437 | **3.09e-27** |
|  | IL-4 | 0.8819201 | **2.87e-20** |
|  | IL-10 | 0.8818740 | **2.90e-20** |
|  | IL-5 | 0.8791033 | **5.40e-20** |
|  | IL-6 | 0.8143054 | **4.38e-15** |
|  | IFNgamma | 0.8046411 | **1.62e-14** |
|  | IL-1beta | 0.7443617 | **1.41e-11** |
|  | CXCL-1 | 0.4578589 | **2.66e-04** |
| **Variance explained: 68.74%** |  |  |  |
| **PCA2** | **Cytokine (hippocampus)** | **Correlation** | **p-value** |
|  | CXCL-1 | 0.5457298 | **7.82e-06** |
|  | IFNgamma | 0.3938848 | **2.02e-03** |
|  | IL-6 | 0.3690473 | **4.02e-03** |
|  | IL-1beta | 0.3380654 | **8.82e-03** |
|  | IL-4 | -0.3296997 | **1.08e-02** |
|  | IL-5 | -0.3514059 | **6.35e-03** |
|  | IL-10 | -0.3742493 | **3.50e-03** |
| **Variance explained: 12.24%** |  |  |  |

For PCA2 IL-1β, IL-6, IFNγ and CXCL-1 loaded positively, whereas IL-4, IL-5 and IL-10 loaded negatively onto this component, which explained 12.24% of the variance. No significant effects between parity and age groups were demonstrated in PCA2 (*p’s* > 0.191; **Fig. 5K**).

For plasma inflammatory signaling (**Fig. 6**) we found that plasma levels of IL-4, IL-6, IL-10 IL-13 and IFNγ were reduced with age (main effect of age: F_(1, 53)_ = 11.990, *p* = 0.001, Ƞ_p_^2^ = 0.184 (IL-4); F_(1, 53)_ = 5.082, *p* = 0.028, Ƞ_p_^2^ = 0.087 (IL-6); F_(1, 53)_ = 10.013, *p* = 0.003, Ƞ_p_^2^ = 0.159 (IL-10); F_(1, 53)_ = 7.450, *p* = 0.009, Ƞ_p_^2^ = 0.123 (IL-13), F_(1, 53)_ = 6.635, *p* = 0.013, Ƞ_p_^2^ = 0.111 (IFNγ); **Fig. 6B, D-G**)). There was a trend towards a significant main effect of parity for plasma IL-1β (F_(2, 53)_ = 2.748, *p* = 0.073, Ƞ_p_^2^ = 0.094: **Fig. 6A**), with higher levels of IL-1β in primiparous compared to nulliparous and biparous rats (primiparous vs nulliparous: *p* = 0.082; primiparous vs biparous: *p* = 0.032). Other than IL-1β, there were no significant main effects of parity or significant interactions between parity and age (all *p*’s > 0.247). A priori comparisons indicated that the age-related reductions in IL-4 and IL-10 were mainly driven by nulliparous and biparous females (IL-4: *p* = 0.005, Cohen’s *d* = 1.229 (nulliparous) and *p* = 0.040, Cohen’s *d* = 0.874 (biparous); IL-10: *p* = 0.025, Cohen’s *d* = 1.044 (nulliparous) and *p* = 0.040, Cohen’s *d* = 0.919 (biparous); **Fig. 6B & E**), while the age-related reduction in IL-6, IL-13 and IFNγ were mainly driven by nulliparous females (IL-6: *p* = 0.001, Cohen’s *d* = 1.010; IL-13: *p* = 0.003, Cohen’s *d* = 0.852; IFNγ: *p* = 0.006, Cohen’s *d* =0.323; **Fig. 6D, F & G)**.

**Figure 6.**
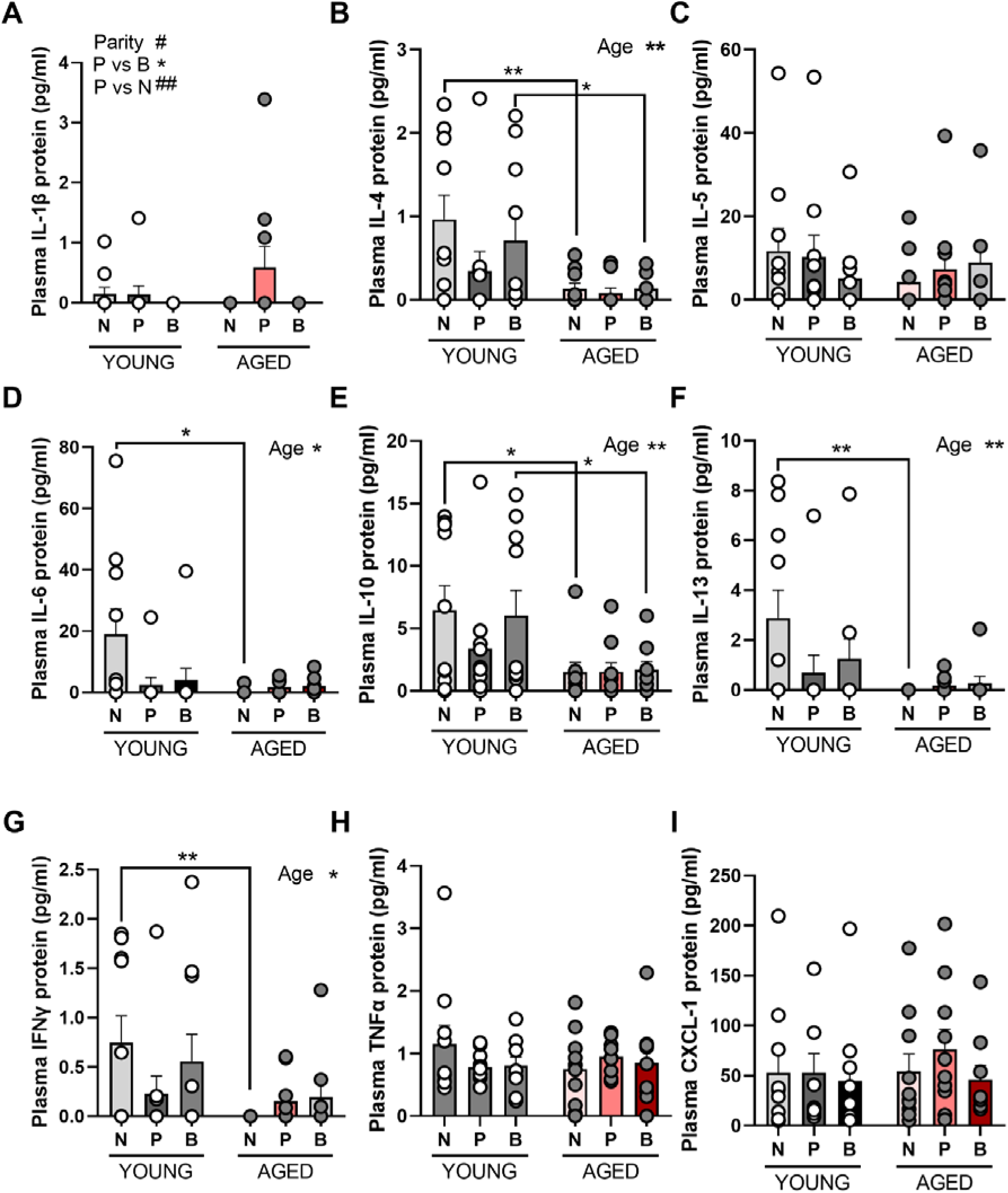
Inflammatory cytokine levels in plasma. Plasma levels of IL-1β varied by parity (A) and plasma levels of IL-4 (B), IL-6 (D), IL-10 (E), IL-13 (F) and IFNγ (G) were reduced with age. No significant differences were found in plasma levels of IL-5 (C), TNFα and CXCL-1 (I) with parity or age . Data are expressed as mean ±SEM. n = 10 (YOUNG) and 9-10 (AGED) per group. * p < 0.05, # *p* = 0.073, ## *p* <u>= 0.082</u>.

**Figure 7.**
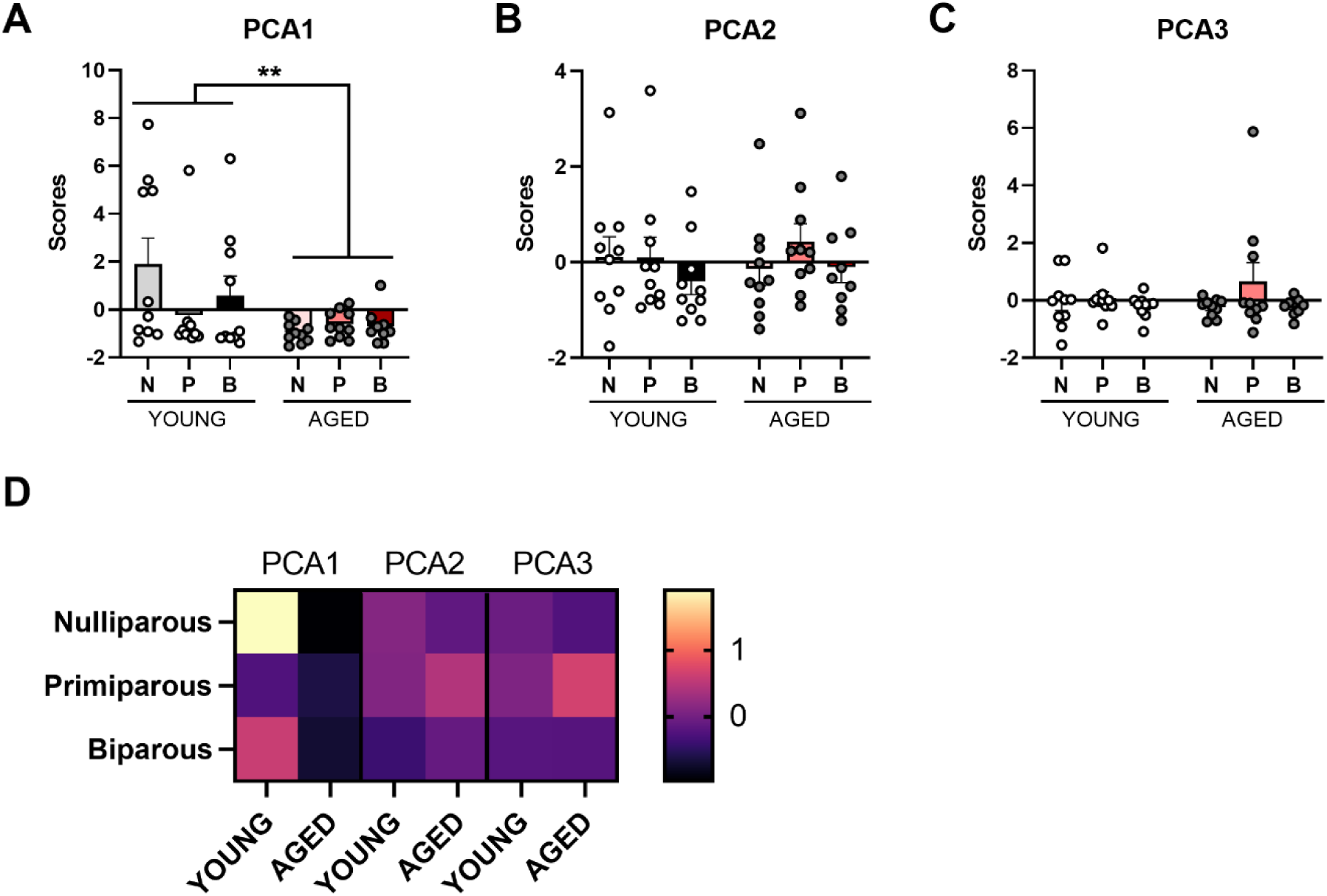
Principal component analysis (PCA) for plasma cytokines. Loadings of scores for each subject onto PCA 1 (A), 2 (B) and 3 (C). Heat map displaying average loading of each parity group on each compartment (D). Data are expressed as mean ±SEM. n = 9-10 (young) and 10 (aged) per group. ** p < 0.01. N = nulliparous, P = primiparous, B = biparous.

### 3.1 Parity transiently alters the tryptophan-kynurenine pathway

The TRP-KYN pathway is stimulated by pro-inflammatory cytokines and is associated with several neurological diseases, such as neurodegenerative disease (Davis and Liu, 2015). Tryptophan, KYN, XA and 3-HAA, were significantly reduced with age (main effect of age: F_(1, 45)_ = 7.298, *p* = 0.010, Ƞ_p_ = 0.140; **Fig. 8A** (tryptophan); F_(1, 43)_ = 20.661, *p* = 0.0004, Ƞ_p_ = 0.325; **Fig. 8B** (KYN); F_(1, 45)_ = 18.061, *p* = 0.0001, Ƞ_p_ = 0.286; **Fig. 8E** (3-HAA); F_(1, 45)_ = 4.567, *p* = 0.038, Ƞ_p_ = 0.092; **Fig. 8G** (XA)).

**Figure 8.**
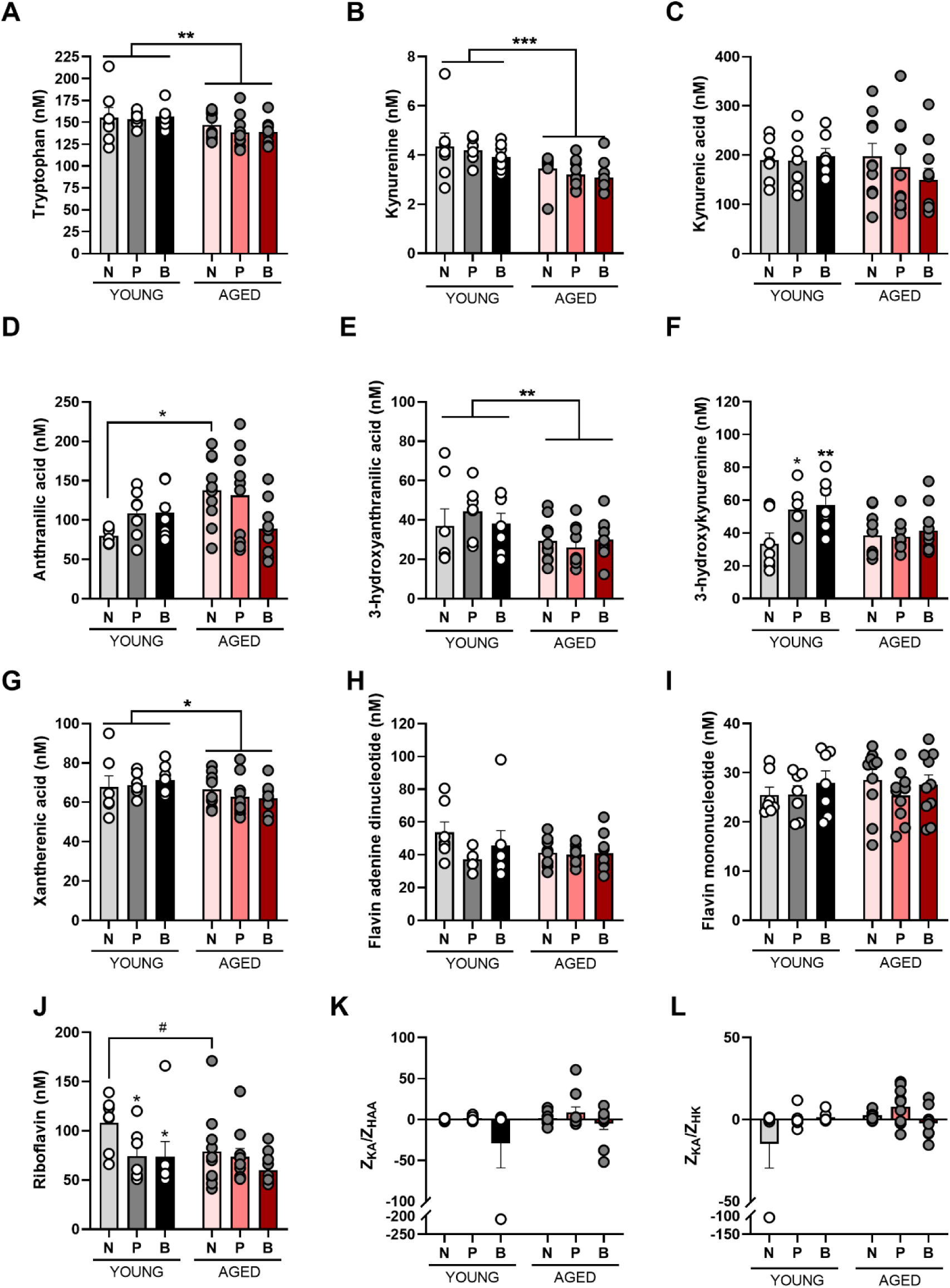
Parity alters the tryptophan-kynurenine pathway in young adult, but not aged females. Plasma levels of tryptophan (A), kynurenine (B), kynurenic acid (C), anthranilic acid (D), 3-hydroxyanthranilic acid (E), 3-hydroxykynurenine (F), xantherenic acid (G), flavin adenine dinucleotide (H), flavin mononucleotide (I) and riboflavin (J). Ratios of Z-scores for neuroprotective plasma metabolite kynurenic acid (KA) and neurotoxic 3-hydroxy anthranilic acid (HAA) and 3-hydroxykynurenine (HK) are demonstrated in K and L, respectively. Data are presented in nM and are expressed as mean ±SEM. n = 6-7 (young) and 9-10 (aged) per group. * p < 0.05, ** p < 0.01, *** p < 0.001, **** p < 0.0001, # *p* = 0.054.

For 3-HK there was a significant interaction between parity and age (F_(2, 45)_ = 3.213, *p* = 0.050, Ƞ_p_ = 0.125; **Fig. 8F**). Post-hoc analysis revealed that young adult nulliparous females had lower levels of 3-HK than both young primiparous (*p* = 0.030, Cohen’s *d* = 1.337) and biparous (*p* = 0.014, Cohen’s *d* = 1.457) females, but this difference was no longer present at middle age (*p* > 0.660).

For the metabolite AA, there was a significant interaction between parity and age (F_(2, 45)_ = 4.143, *p* = 0.022, Ƞ_p_^2^ = 0.155; **Fig. 8D**). Post hoc analysis revealed that nulliparous aged females had significantly higher levels of AA than young adult nulliparous females (*p* = 0.045, Cohen’s *d* = 1.897).

Next, we investigated the levels of the enzyme FAD, which promotes the neurotoxic arm in the TRP-KYN pathway. There were no significant differences in FAD levels between age or parity groups (all *p’s* > 0.140; **Fig. 8H**). FMN was also not affected by age and/or parity (*p*’s > 0.526; **Fig. 8I**).

Several key enzymes in the TRP-KYN pathway also require riboflavin (vitamin B2) as a cofactor (Christensen et al., 2018), and riboflavin is an important vitamin during pregnancy that’s heavily utilized during both pregnancy and postpartum. Young adult nulliparous females had significantly higher levels of riboflavin than the two young adult parous groups (primiparity: *p* = 0.041, Cohen’s *d* = 1.272; biparity: *p* = 0.037, Cohen’s *d* = 0.999, but there were no significant differences in the middle-aged group (parity by age interaction effect: F_(2, 42)_ = 4.579, *p* = 0.016, Ƞ_p_ = 0.179; **Fig. 8J**)). There were also significant main effects of parity (F_(2, 42)_ = 8.528, *p* = 0.0007, Ƞ_p_ = 0.289) and age (F_(1, 42)_ = 6.848, *p* = 0.012, Ƞ_p_ = 0.140).

Assessing a neuroprotective vs neurotoxic ratio of KA and 3-HAA, factors of the TRP-KYN pathway, did not reveal any significant main effects of parity, age, nor an interaction between the two (all *p’s* > 0.101; **Fig. 8K**). Neither were there differences between groups when assessing the ratio of KA and HL (all *p’s* > 0.426; **Fig. 8L**).

We also analyzed the TRP-KYN metabolites using PCA. Component loading are displayed in Table 3. Results from the PCA analysis yielded three principal components, which together explained 81.11% of the total variance. TRP, XA, KYN, 3-HK, 3-HAA and KA loaded positively onto principal component analysis (PCA) 1 which explained 49.97% of the variance (**Fig. 9A**). In PCA1, middle-aged females had lower scores than young adult females (main effect of age: F_(1, 45)_ = 10.622, *p* = 0.002, Ƞ_p_ = 0.191) but no other significant effects ( *p’s* > 0.379).

**Figure 9.**
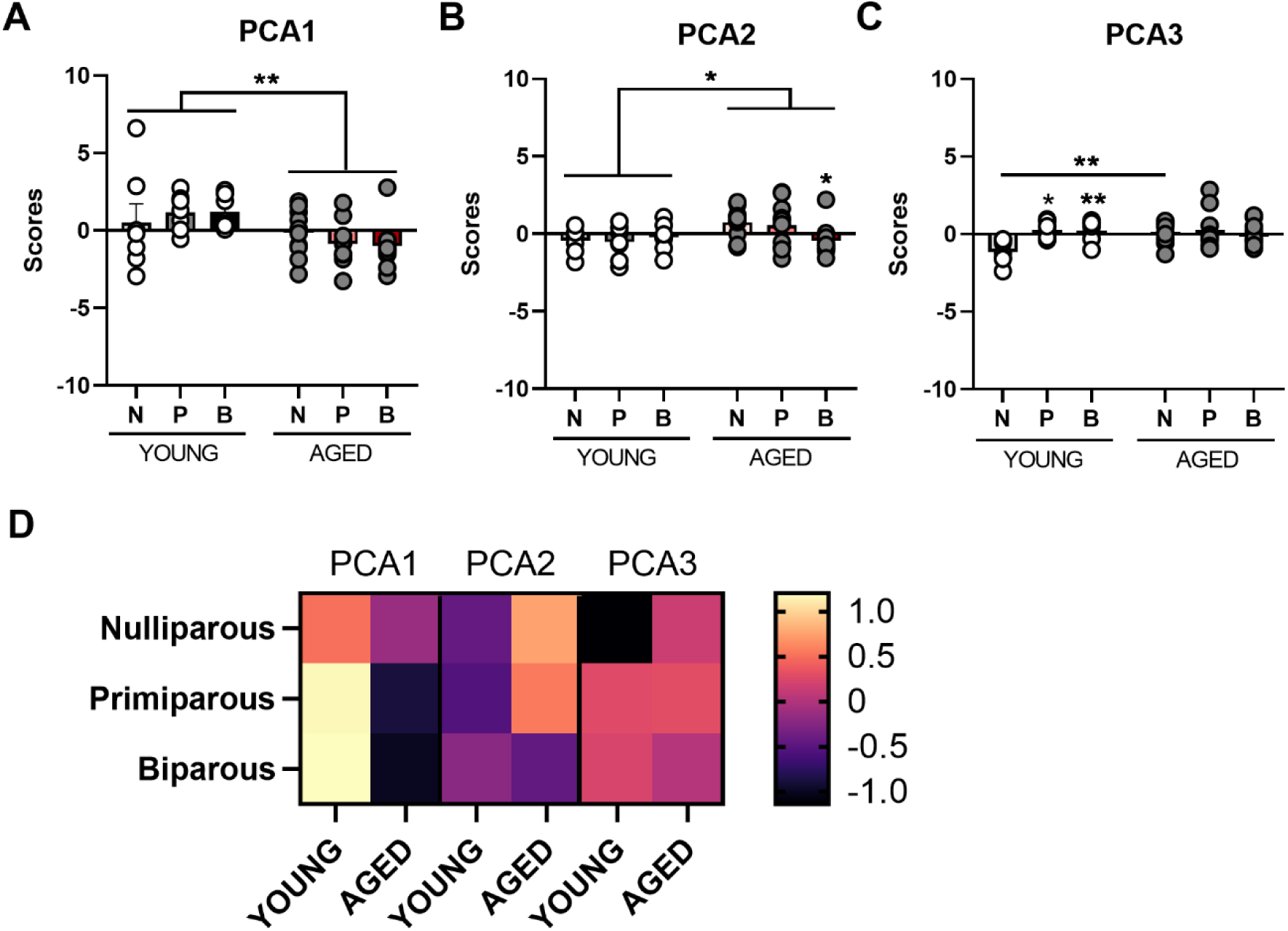
Principal component analysis (PCA) for tryptophan-kynurenine metabolites. Loadings of scores for each subject onto PCA 1 (A), 2 (B) and 3 (C). Heat map displaying average loading of each parity group on each compartment (D). Data are expressed as mean ±SEM. n = 7 (young) and 10 (aged) per group. * p < 0.05, ** p < 0.01. N = nulliparous, P = primiparous, B = biparous.

**Table 2.**
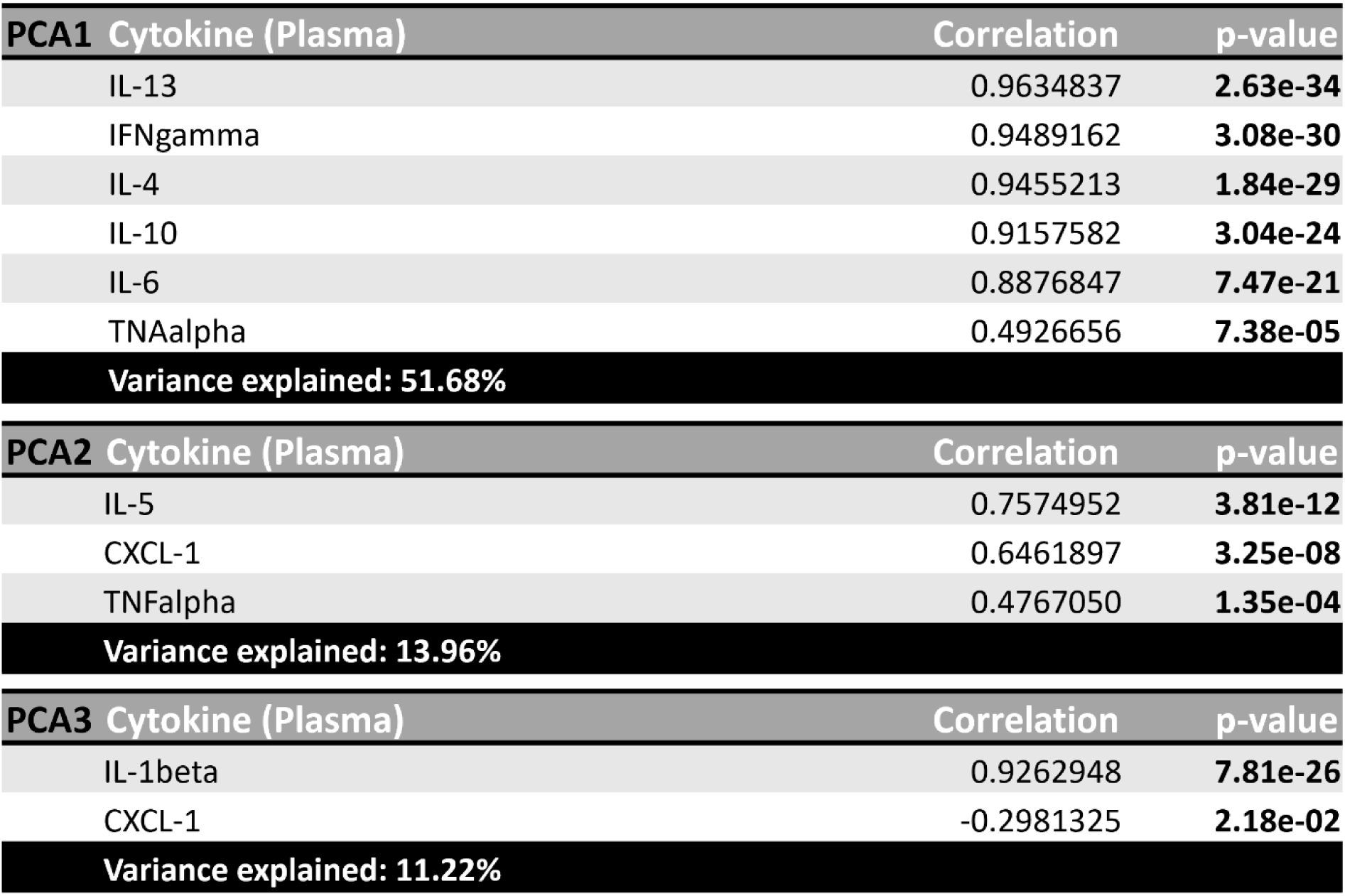
Component loadings of the cytokine variance, in plasma, of components identified using principal component analysis (PCA).

**Table 3.** Component loadings of the plasma metabolite variance of components identified using principal component analysis (PCA).

| PCA1 | Metabolic factor | Correlation | p-value |
| --- | --- | --- | --- |
|  | Tryptophan | 0.923276 | **5.34E-22** |
|  | Xanthurenic acid | 0.9222197 | **7.38E-22** |
|  | Kynurenine | 0.8381485 | **1.69E-14** |
|  | 3-hydroxykynurenine | 0.6823298 | **3.50E-08** |
|  | 3-hydroxyanthranilic acid | 0.6238258 | **1.01E-06** |
|  | Kynurenic acid | 0.4251385 | **1.87E-03** |
| **Variance explained: 49.97%** |  |  |  |
| PCA2 | Metabolic factor | Correlation | p-value |
|  | Anthranilic acid | 0.7237361 | **1.97e-09** |
|  | Kynurenic acid | 0.6562456 | **1.71e-07** |
|  | 3-hydroxyanthranilic acid | -0.5331410 | **5.62e-05** |
| **Variance explained: 19.49%** |  |  |  |
| PCA3 | Metabolic factor | Correlation | p-value |
|  | Anthranilic acid | 0.5492708 | **2.99e-05** |
|  | 3-hydroxykynurenine | 0.5209544 | **8.88e-05** |
| **Variance explained: 11.65%** |  |  |  |

For PCA2, AA and KA loaded positively, whereas 3-HAA loaded negatively onto this component, which explained 19.49% of the variance. Age resulted in higher scores at middle age, compared to in younger females (main effect of age: F_(1, 45)_ = 4.735, *p* = 0.035, Ƞ_p_ = 0.095; **Fig. 9B**). There were no other significant effects (*p’s >* 0.128).

Lastly, AA and 3-HK, loaded positively to PCA3 which explained 11.65% of the variance (**Fig. 9C**). In PCA3 there is a significant interaction between parity and age (interaction: F_(2, 45)_ = 3.849, *p* = 0.029, Ƞ_p_^2^ = 0.146). Post-hoc analysis revealed that nulliparous females displayed significantly lower scores than primiparous (*p* = 0.010, Cohen’s *d* = 2.133) and biparous (*p* = 0.009, Cohen’s *d* = 1.866) females in the young adult age group. In addition, young adult nulliparous females had lower scores than middle age nulliparous females (*p* = 0.008, Cohen’s *d* = 1.733) but this was not seen in the parous groups (*p’s* > 0.878). There was no significant effect of age (*p* = 0.124).

### 3.5 Vitamin B6 levels are reduced with age

The TRP-KYN pathway is sensitive to vitamin B6 availability, in which two of the key enzymes kynurenine aminotransferase and kynureninase require the active form of B6, pyridoxal 5’phosphate. Vitamin B6 is required for many processes during pregnancy, and vitamin B6 deficiency is common during this period (van de Kamp and Smolen, 1995). Therefore, we measured vitamin B6 and its active coenzyme forms. The levels of pyridoxic acid, pyridoxal, pyridoxal 5’-phosphate and pyridoxamine were significantly reduced with age (main effect of age: F_(1, 45)_ = 7.768, *p* = 0.008, Ƞ_p_^2^ = 0.147, **Fig. 10A** (pyridoxic acid); F_(1, 43)_ = 6.563, *p* = 0.014, Ƞ_p_^2^ = 0.132, **Fig. 10B** (pyridoxal); F_(1, 45)_ = 6.619, *p* = 0.013, Ƞ_p_^2^ = 0.128, **Fig. 10C** (pyridoxal 5’-phosphate); F_(1, 45)_ = 4.710, *p* = 0.035, Ƞ_p_^2^ = 0.095, **Fig. 10D** (pyridoxamine)), but there were no other significant main or interaction effects ( *p’s >* 0.235). For pyridoxine, neither the main effect of parity, age, nor the interaction were significant ( *p* > 0.135, **Fig. 10E**).

**Figure 10.**
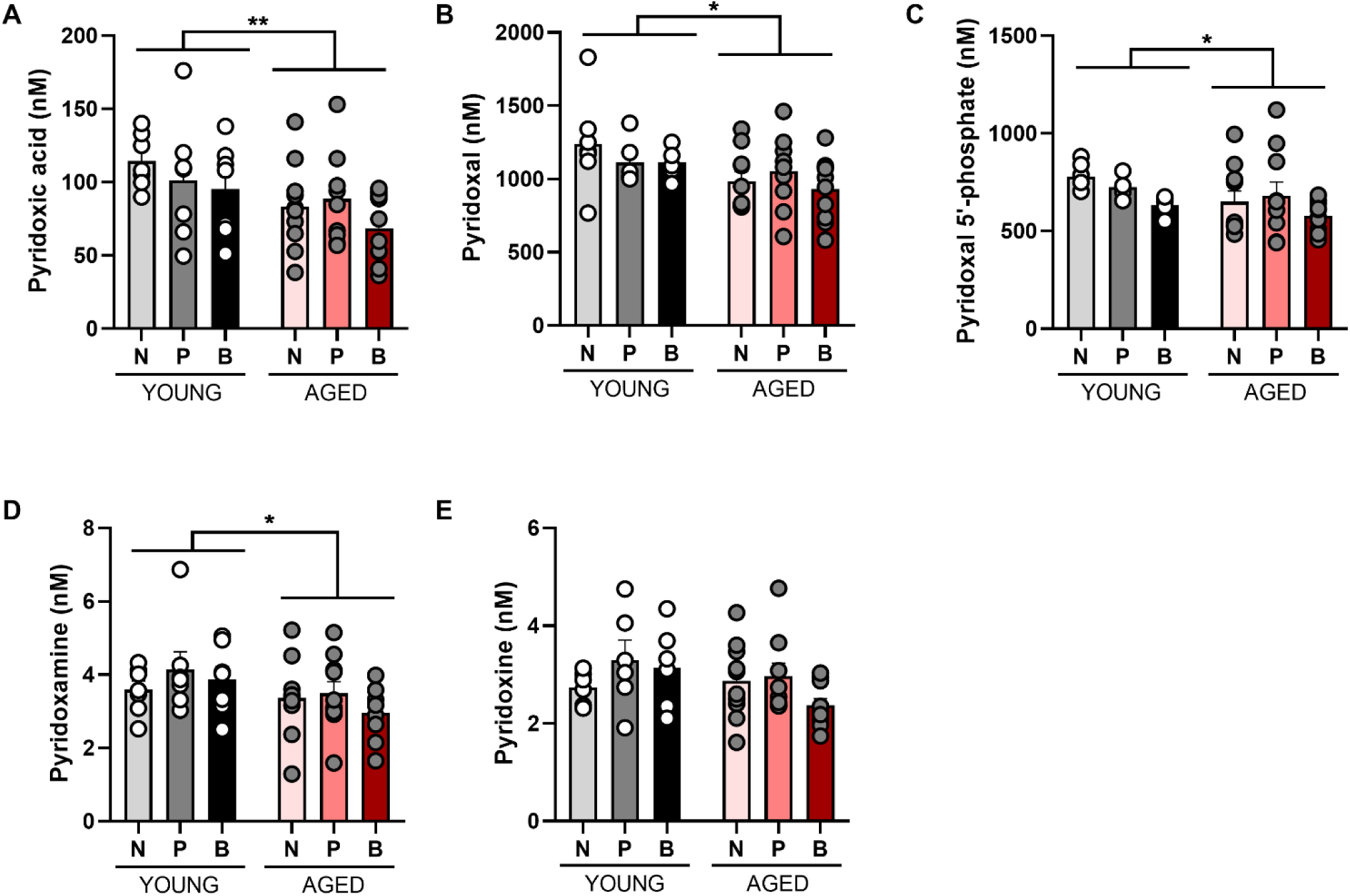
Levels of vitamin B6 coenzymes in plasma are reduced with age, regardless of parity. Levels of vitamin B6 coenzymes pyridoxic acid (A), pyridoxal (B), pyridoxal 5’-phostphate (C) and pyridoxamine (D) were reduced with age, but not or pyridoxine (E). Data are presented in nM and expressed as mean ±SEM. n = 7 (young) and 10 (aged) per group. * p < 0.05, ** p < 0.01.

## 4 Discussion

Pregnancy and the postpartum involves vast physiological changes, including effects on the central nervous system. Here we conduct a broad survey of cellular and molecular signatures of motherhood in the adult and ageing hippocampus. We examined the effects of primiparity and biparity on central and peripheral inflammation, metabolomics, and synaptic plasticity after parturition and weaning in the short term (postpartum day 30) and at middle age (postpartum day 240). We found that parity results in short and long-term changes in the synaptic protein PSD-95, cell signalling, inflammatory signalling, and inflammation-associated TRP-KYN pathway. The amount of parity was also important for certain outcomes as biparity had greater effects on inflammatory-related signalling (cell signalling proteins, and microglia). Lastly, parity prevented the age-related decline in stem cells observed in nulliparous rats, suggesting that parity alters the trajectory of brain aging.

### 4.1 Parity regulated aspects of neuroplasticity in the short and in the long-term

In the current study we found that previous parity increased levels of PSD-95, pp70S6K and S6RP, effects that were detected at postpartum day 30 and lasted until middle-age. Notably, effects on cell signalling phosphoproteins were only seen in biparous animals, highlighting the importance of considering amount of parity. PSD-95 is a very abundant scaffold protein in the PSD fraction and its role is to organize the PSD by anchoring mostly glutamate channels and receptors (Hunt et al., 1996; Kim and Sheng, 2004). Both phosphoproteins that were increased with biparity play important roles in synaptic plasticity. Furthermore, elevated S6RP phosphorylation is associated with increased synaptic plasticity in hippocampal, cortical, and striatal neurons (Antion et al., 2008; Bowling et al., 2014; Kelleher et al., 2004; Klann and Dever, 2004; Tsokas et al., 2007, 2005). Pp70S6K, a protein in the protein kinase B (Akt) pathway, plays an important role in synaptic plasticity (Hers et al., 2011). Thus, overall, we find that proteins related to synaptic plasticity are upregulated with parity in the hippocampus.

Our findings are consistent with previous work demonstrating increases in the synaptic proteins, spinophilin and synaptophysin, in the hippocampus or whole brain of multiparous rodents (mice and rats) in middle age (Cui et al., 2014; Rossetti et al., 2016). Parity increases dendritic spine density in the CA1 region in late pregnancy, early postpartum (in lactating females; 5-6 days postpartum) and in older age (18 month) (Flores-Vivaldo et al., 2019; Kinsley et al., 2006). However, primiparity reduces CA3 and CA1 dendritic number and length in the late postpartum compared to biparous or nulliparous female rats (Pawluski and Galea, 2007). In middle and older age, previous parity increases CA1 long-term potentiation in rodents (Lemaire et al., 2006). Together with the results of the current study, this indicates that increases in synaptic proteins are a feature of motherhood that is related to the amount of reproductive experience (primiparous, biparous, multiparous). Although functional implication are not clear, studies indicate that synaptic protein levels are associated with cognitive ability in males and females (Frick and Fernandez, 2003; Ning et al., 2020). Our findings are therefore consistent with previous studies reporting improvements in spatial learning and memory in parous females at the same time points (30 days postpartum and middle age) (Barha et al., 2015; Cui et al., 2014; Gatewood et al., 2005; Macbeth et al., 2008; Pawluski et al., 2006). Maternal changes to synaptogenesis and cognition are similar to environmental enrichment in rodents, suggesting that the maternal experience is stimulating and protective for brain health (Orchard et al., 2023)In humans, de Lange and colleagues demonstrated that a higher number of childbirths was associated with less apparent brain aging, including in the hippocampus (de Lange et al., 2020, 2019; Lange et al., 2020). Thus, parity is associated with increases in synaptic proteins which may indicate improved memory and reduced brain aging, although more research is necessary to establish this connection.

### 4.2 Biparity was associated with greater density of microglia in the dentate gyrus in females

In the present study, biparity resulted in a lasting increase in Iba1+ cell density in the dentate gyrus, observed regardless of age when compared to nulliparous females. Although studies investigating age-related changes in microglia have produced varying results, this may be due to inconsistent inclusion of hippocampal subregions (Lynch, 2022), sex, and parity (due to the use of retired breeders). One previous study including females found an age-related increase in microglia in the dentate gyrus which was not present between young and aged males (Mouton et al., 2002). An age-dependent increase in Iba1+ density was also reported in the dentate gyrus of primiparous, but not nulliparous, females (Eid et al., 2019).

Biparous females also showed a trend towards a reduction in hippocampal levels of CXCL-1 with age. CXCL-1 is an important chemokine for neutrophil recruitment. Hippocampal cytokines remained largely unaltered by parity and age. It is important to note that we measured cytokines in whole hippocampal samples, therefore it is possible that subregional differences were missed, for instance in the dentate gyrus which had the greatest density of Iba1+ cells. Primiparity was associated with an increase in plasma IL-1β with age, unlike in nulliparous or biparous rats, suggesting a possible long-lasting proinflammatory signalling with parity. However most other proinflammatory cytokines were not affected by parity, but were reduced with age. We did not find any differences in the levels of CXCL-1 in plasma, consistent with previous results from our lab (Eid et al., 2019), albeit earlier work only included nulliparous and primiparous females. Future work should consider other areas of the brain and regional differences.

### 4.3 Parity prevented the age-related decline in neural stem cells

Neural stem cells were reduced with age, as expected (Lecanu et al., 2010), but only in the nulliparous group. The same aging effects were not seen in parous groups, indicating that previous parity may mitigate age-dependent declines in neural stem cells. These findings may explain the greater levels of neurogenesis in the hippocampus of parous rats in middle-age compared to nulliparous rats (Barha et al., 2015; Eid et al., 2019; Galea et al., 2018). Further studies should be conducted to determine whether these neural stem cells are more likely to be dividing in middle aged parous compared to nulliparous rats. These results suggest that previous parity may alter the trajectory of brain aging, particularly in the hippocampus.

### 4.4 Parity transiently alters the tryptophan-kynurenine pathway

The TRP-KYN pathway interacts with the immune system and its enzymes are expressed in many immune cells, including microglia and macrophages (Guillemin et al., 2003). Only two molecules differed between parity groups postpartum: 3-HK and riboflavin, also known as vitamin B2. 3-HK is increased during pregnancy (Ceresoli-Borroni and Schwarcz, 2000), and we found levels of this metabolite remained significantly elevated in both primiparous and biparous females, but there were no differences between parous groups in middle age. Vitamin B2 levels were significantly lower in primiparous and biparous females at 30 days postpartum, compared to age-matched nulliparous females, but these differences were no longer present at middle age. The risk of vitamin B2 deficiency is highest during the third trimester and lactation (Plantone et al., 2021) and our results suggest that lower levels can persist in rats at least a week after the pups have been weaned. Although KYN, KA and XA are commonly elevated during pregnancy, we found no differences in the levels of these kynurenines at 30 days postpartum, indicating that their levels were not affected by the reduction of riboflavin in parous females. Not surprisingly, the most prominent changes to TRP-KYN metabolite levels were by age, as levels of TRP, 3-HAA and XA were reduced at middle age, compared to young adult females, consistent with other findings (Comai et al., 2005; Frick et al., 2004; Pertovaara et al., 2006). Reductions in TRP and XA have previously been linked to an increased risk of cognitive impairment (Bakker et al., 2021). However, in contrast to our results, KYN has previously been found to increase with age (Braidy et al., 2011; de Bie et al., 2016).

Levels of AA, the third immediate downstream KYN metabolite, were increased with age, but only in nulliparous females. In addition, HK was increased 30 days postpartum in primiparous and biparous females, compared to nulliparous females. An increase in this metabolite has also been reported in humans around 40 days postpartum (Veen et al., 2016).

Pyridoxal 5’ phosphate, an active form of vitamin B6, reduces hippocampal apoptosis by down-regulating inflammation, up-regulating brain-derived neurotrophic factor and preventing the exhaustion of cellular energy stores in rats (sex was not reported)(Zysset-Burri et al., 2013). Lower levels of plasma pyridoxal 5’ phosphate have previously been demonstrated in older adults (Bates et al., 1999; Morris et al., 2008) and pyridoxine deficiency is associated with cognitive impairment in wildtype male mice and in animal models of Alzheimer’s disease (Hasegawa et al., 2010; Jung et al., 2020). In line with this, we found reduced levels of pyridoxal 5’-phosphate with age. Age-related reductions were also found for other forms of vitamin B6, including pyridoxic acid, pyridoxal, and pyridoxamine. However, parity did not alter the levels of any form of vitamin B6 postpartum or at middle age.

### 4.5 Parity alters the trajectory of aging

Intriguingly, some effects of parity were only noted in the biparous group. Biparity increased levels of cell signalling phosphoproteins, metabolites, a chemokine, and microglia in the dentate gyrus. Other groups have found that the amount of parity influences cognition, neuroplasticity and potentially risk for aging-related disorders (de Lange et al., 2020; Glynn, 2012; Pawluski and Galea, 2007). For example, biparity was associated with greater survival of new neurons across the postpartum (Pawluski and Galea, 2007). Other findings suggest that while the levels of certain proteins were not different in middle age, the aging trajectory was altered with parity. Here, we report that aging decreased neural stem cells in the dentate gyrus, and AA, 3-HK and riboflavin in plasma of nulliparous but not parous rats. This may be related to findings indicating less evident brain aging in humans with childbirths, in the hippocampus as well as other regions (Lange et al., 2020). Lastly, Orchard and colleagues proposed that reproductive experience in humans may result in similar neural and cognitive benefits as environmental enrichment in rodents (Orchard et al., 2023). More specifically, the continual adjustments and changing environments that are needed to adapt to during a child’s development result in sensory, cognitive, motor and social stimulation in the mother (Orchard et al., 2023). These effects can stimulate synaptogenesis (Diamond, 1988; Diamond et al., 1964) and may be protective against brain aging (Speisman et al., 2013).

## 5 Conclusions

We found that on a variety of neuroplastic and inflammatory markers, parity has a significant impact long after parturition, including some effects that lasted into middle-age. Given that parity is associated with different disease risk for stroke, diabetes, and Alzheimer’s disease (Huo et al., 2021; Lee et al., 2022; Parikh et al., 2010), it is important to understand how the brain may change with parity and age. In our study we found that parity increased PSD-95 and microglia levels in the hippocampus. We also found that parity altered the trajectory of aging, as although neural stem cell density decreased in the hippocampus of nulliparous rats, this same profile was not seen in parous animals with age. Although biparous mothers had a higher level of microglia in the dentate gyrus than nulliparous females we did not see any robust parity-based changes in circulating or central inflammatory cytokine levels. Future work could examine a more regional approach to inflammatory cytokines. The TRP-KYN system was mainly affected by age, with a reduction in the levels of several key metabolites of this pathway at middle age. Although some parity related-changes were found in TRP-KYN metabolites, these were limited to young adult females and did not persist until middle age. Therefore, parity alters the trajectory of brain aging for certain parameters, whereas others were affected specifically by the presence of previous parity or age, solely. Further research is necessary to investigate if these changes may contribute to disease under certain conditions.

## 6 Supplementary material

**Supplementary Figure 1.**
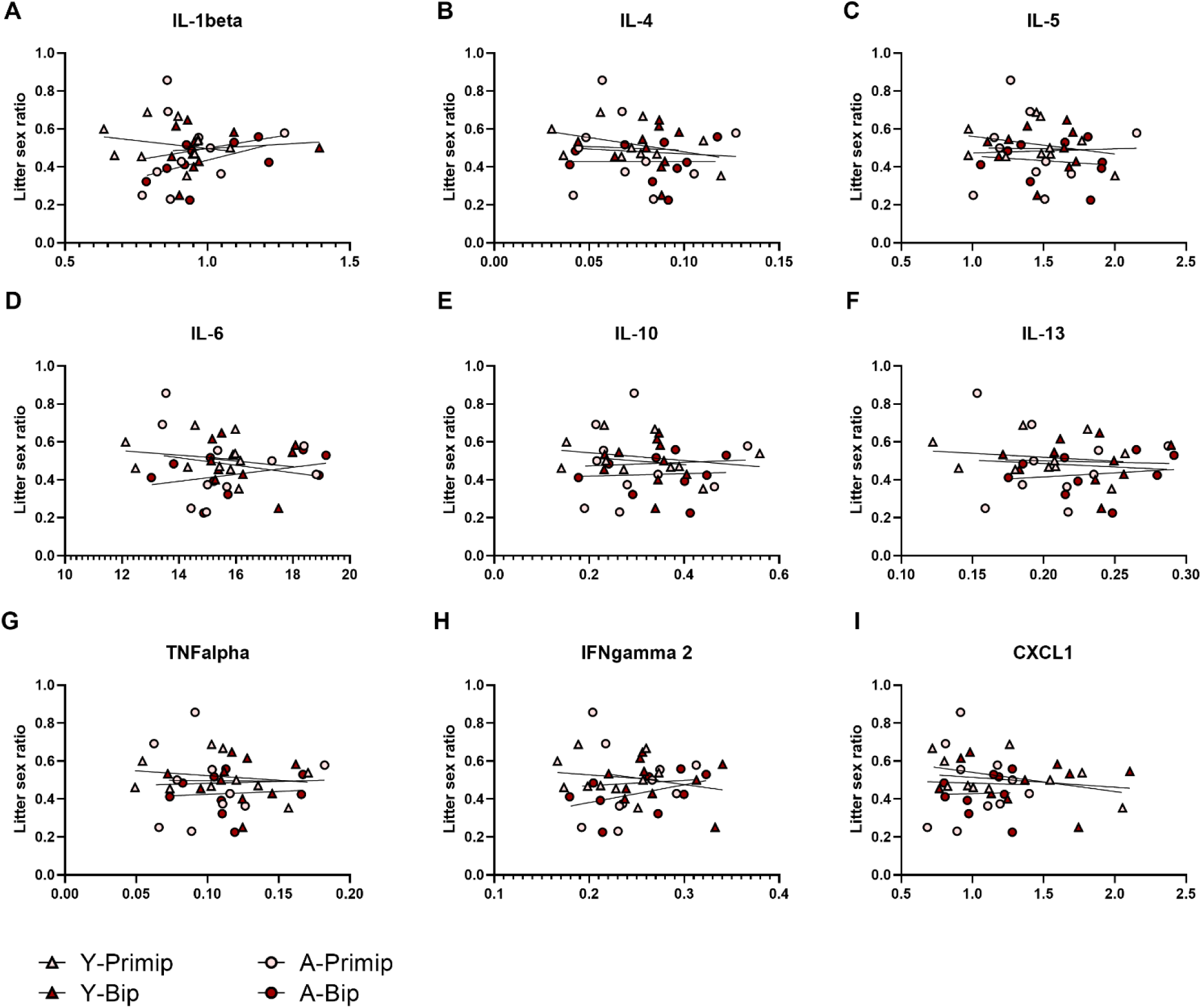
Cytokine levels were not influenced by the ratio of males and females in each litter. Sex litter ratio did not significantly effect cytokine levels of (IL-1β (A), IL-4 (B), IL-5 (C), IL-6 (D), IL-10 (E), IL-13 (F), TNFα (G), IFNγ (H) or CXCL-1 (I). n = 10 (YOUNG) and 9-10 (AGED) per group.

## 7 Acknowledgements

We gratefully acknowledge Kimberly Go, Nicole Minielly, Jessica Chaiton, and Muna Ibrahim for help with breeding and animal care and the animal care staff at Centre for Disease Modelling at University of British Columbia.

## 9 Author Contributions

Paula Duarte-Guterman: Conceptualization, Methodology, Data curation, Investigation, writing original draft; Funding acquisition. Jennifer Richard: Data curation, Visualization, Formal analysis, Writing-Original draft preparation. Rand Eid: Investigation, Writing-review and editing. Stephanie Lieblich: Methodology, Investigation, Writing Review/editing. Yvonne Lamers: Investigation. Liisa Galea: Conceptualization, Formal Analysis, Funding acquisition, Supervision, Writing-Reviewing and Editing.

## References

Aagaard-Tillery, K.M., Silver, R., Dalton, J., 2006. Immunology of normal pregnancy. Seminars in Fetal and Neonatal Medicine 11, 279–295. https://doi.org/10.1016/j.siny.2006.04.003

Abbassi-Ghanavati, M., Greer, L.G., Cunningham, F.G., 2009. Pregnancy and laboratory studies: a reference table for clinicians. Obstet Gynecol 114, 1326–1331. https://doi.org/10.1097/AOG.0b013e3181c2bde8

Amrein, I., 2015. Adult Hippocampal Neurogenesis in Natural Populations of Mammals. Cold Spring Harb Perspect Biol 7, a021295. https://doi.org/10.1101/cshperspect.a021295

Antion, M.D., Hou, L., Wong, H., Hoeffer, C.A., Klann, E., 2008. mGluR-Dependent Long-Term Depression Is Associated with Increased Phosphorylation of S6 and Synthesis of Elongation Factor 1A but Remains Expressed in S6K-Deficient Mice. Molecular and Cellular Biology 28, 2996–3007. https://doi.org/10.1128/MCB.00201-08

Badawy, A.A.-B., 2015. Tryptophan metabolism, disposition and utilization in pregnancy. Biosci Rep 35, e00261. https://doi.org/10.1042/BSR20150197

Bakker, L., Ramakers, I.H.G.B., van Boxtel, M.P.J., Schram, M.T., Stehouwer, C.D.A., van der Kallen, C.J.H., Dagnelie, P.C., van Greevenbroek, M.M.J., Wesselius, A., Midttun, Ø., Ueland, P.M., Verhey, F.R.J., Eussen, S.J.P.M., Köhler, S., 2021. Associations between plasma kynurenines and cognitive function in individuals with normal glucose metabolism, prediabetes and type 2 diabetes: the Maastricht Study. Diabetologia 64, 2445–2457. https://doi.org/10.1007/s00125-021-05521-4

Barha, C.K., Galea, L.A.M., 2017. The maternal “baby brain” revisited. Nat Neurosci 20, 134–135. https://doi.org/10.1038/nn.4473

Barha, C.K., Lieblich, S.E., Chow, C., Galea, L.A.M., 2015. Multiparity-induced enhancement of hippocampal neurogenesis and spatial memory depends on ovarian hormone status in middle age. Neurobiology of Aging 36, 2391–2405. https://doi.org/10.1016/j.neurobiolaging.2015.04.007

Barth, C., de Lange, A.-M.G., 2020. Towards an understanding of women’s brain aging: the immunology of pregnancy and menopause. Frontiers in Neuroendocrinology 58, 100850. https://doi.org/10.1016/j.yfrne.2020.100850

Bates, C.J., Pentieva, K.D., Prentice, A., Mansoor, M.A., Finch, S., 1999. Plasma pyridoxal phosphate and pyridoxic acid and their relationship to plasma homocysteine in a representative sample of British men and women aged 65 years and over. Br J Nutr 81, 191–201.

Berg, S., Kutra, D., Kroeger, T., Straehle, C.N., Kausler, B.X., Haubold, C., Schiegg, M., Ales, J., Beier, T., Rudy, M., Eren, K., Cervantes, J.I., Xu, B., Beuttenmueller, F., Wolny, A., Zhang, C., Koethe, U., Hamprecht, F.A., Kreshuk, A., 2019. ilastik: interactive machine learning for (bio)image analysis. Nat Methods 16, 1226–1232. https://doi.org/10.1038/s41592-019-0582-9

Bowling, H., Zhang, G., Bhattacharya, A., Pérez-Cuesta, L.M., Deinhardt, K., Hoeffer, C.A., Neubert, T.A., Gan, W., Klann, E., Chao, M.V., 2014. Antipsychotics Activate mTORC1-Dependent Translation to Enhance Neuronal Morphological Complexity. Science Signaling 7, ra4–ra4. https://doi.org/10.1126/scisignal.2004331

Braidy, N., Guillemin, G.J., Mansour, H., Chan-Ling, T., Grant, R., 2011. Changes in kynurenine pathway metabolism in the brain, liver and kidney of aged female Wistar rats: Changes in kynurenine pathway metabolism in aging rats. FEBS Journal 278, 4425–4434. https://doi.org/10.1111/j.1742-4658.2011.08366.x

Carmona, S., Martínez-García, M., Paternina-Die, M., Barba-Müller, E., Wierenga, L.M., Alemán-Gómez, Y., Pretus, C., Marcos-Vidal, L., Beumala, L., Cortizo, R., Pozzobon, C., Picado, M., Lucco, F., García-García, D., Soliva, J.C., Tobeña, A., Peper, J.S., Crone, E.A., Ballesteros, A., Vilarroya, O., Desco, M., Hoekzema, E., 2019. Pregnancy and adolescence entail similar neuroanatomical adaptations: A comparative analysis of cerebral morphometric changes. Hum Brain Mapp 40, 2143–2152. https://doi.org/10.1002/hbm.24513

Castro-Portuguez, R., Sutphin, G.L., 2020. Kynurenine pathway, NAD+ synthesis, and mitochondrial function: Targeting tryptophan metabolism to promote longevity and healthspan. Experimental Gerontology 132, 110841. https://doi.org/10.1016/j.exger.2020.110841

Ceresoli-Borroni, G., Schwarcz, R., 2000. Perinatal kynurenine pathway metabolism in the normal and asphyctic rat brain. Amino Acids 19, 311–323. https://doi.org/10.1007/s007260070062

Chauhan, G., Tadi, P., 2022. Physiology, Postpartum Changes, in: StatPearls. StatPearls Publishing, Treasure Island (FL).

Christensen, M.H.E., Fadnes, D.J., Røst, T.H., Pedersen, E.R., Andersen, J.R., Våge, V., Ulvik, A., Midttun, Ø., Ueland, P.M., Nygård, O.K., Mellgren, G., 2018. Inflammatory markers, the tryptophan-kynurenine pathway, and vitamin B status after bariatric surgery. PLoS One 13, e0192169. https://doi.org/10.1371/journal.pone.0192169

Comai, S., Costa, C.V.L., Ragazzi, E., Bertazzo, A., Allegri, G., 2005. The effect of age on the enzyme activities of tryptophan metabolism along the kynurenine pathway in rats. Clinica Chimica Acta 360, 67–80. https://doi.org/10.1016/j.cccn.2005.04.013

Cui, J., Jothishankar, B., He, P., Staufenbiel, M., Shen, Y., Li, R., 2014. Amyloid precursor protein mutation disrupts reproductive experience-enhanced normal cognitive development in a mouse model of Alzheimer’s disease. Mol Neurobiol 49, 103–112. https://doi.org/10.1007/s12035-013-8503-x

Davis, I., Liu, A., 2015. What is the tryptophan kynurenine pathway and why is it important to neurotherapy? Expert Rev Neurother 15, 719–721. https://doi.org/10.1586/14737175.2015.1049999

de Bie, J., Guest, J., Guillemin, G.J., Grant, R., 2016. Central kynurenine pathway shift with age in women. J. Neurochem. 136, 995–1003. https://doi.org/10.1111/jnc.13496

de Lange, A.-M.G., Barth, C., Kaufmann, T., Maximov, I.I., van der Meer, D., Agartz, I., Westlye, L.T., 2020. Women’s brain aging: Effects of sex-hormone exposure, pregnancies, and genetic risk for Alzheimer’s disease. Hum Brain Mapp 41, 5141–5150. https://doi.org/10.1002/hbm.25180

de Lange, A.-M.G., Kaufmann, T., van der Meer, D., Maglanoc, L.A., Alnæs, D., Moberget, T., Douaud, G., Andreassen, O.A., Westlye, L.T., 2019. Population-based neuroimaging reveals traces of childbirth in the maternal brain. Proc Natl Acad Sci U S A 116, 22341–22346. https://doi.org/10.1073/pnas.1910666116

Diamond, M.C., 1988. Enriching heredity: The impact of the environment on the anatomy of the brain, Enriching heredity: The impact of the environment on the anatomy of the brain. Free Press, New York, NY, US.

Diamond, M.C., Krech, D., Rosenzweig, M.R., 1964. The effects of an enriched environment on the histology of the rat cerebral cortex. Journal of Comparative Neurology 123, 111–119. https://doi.org/10.1002/cne.901230110

Duarte-Guterman, P., Leuner, B., Galea, L.A.M., 2019. The long and short term effects of motherhood on the brain. Frontiers in Neuroendocrinology 53, 100740. https://doi.org/10.1016/j.yfrne.2019.02.004

Eid, R.S., Chaiton, J.A., Lieblich, S.E., Bodnar, T.S., Weinberg, J., Galea, L.A.M., 2019. Early and late effects of maternal experience on hippocampal neurogenesis, microglia, and the circulating cytokine milieu. Neurobiology of Aging 78, 1–17. https://doi.org/10.1016/j.neurobiolaging.2019.01.021

Facca, T.A., Mastroianni-Kirsztajn, G., Sabino, A.R.P., Passos, M.T., dos Santos, L.F., Famá, E.A.B., Nishida, S.K., Sass, N., 2018. Pregnancy as an early stress test for cardiovascular and kidney disease diagnosis. Pregnancy Hypertension 12, 169–173. https://doi.org/10.1016/j.preghy.2017.11.008

Flores-Vivaldo, Y.M., Camacho-Abrego, I., Picazo, O., Flores, G., 2019. Pregnancies alters spine number in cortical and subcortical limbic brain regions of old rats. Synapse 73, e22100. https://doi.org/10.1002/syn.22100

Frick, B., Schroecksnadel, K., Neurauter, G., Leblhuber, F., Fuchs, D., 2004. Increasing production of homocysteine and neopterin and degradation of tryptophan with older age. Clinical Biochemistry 37, 684–687. https://doi.org/10.1016/j.clinbiochem.2004.02.007

Frick, K.M., Fernandez, S.M., 2003. Enrichment enhances spatial memory and increases synaptophysin levels in aged female mice. Neurobiol Aging 24, 615–626. https://doi.org/10.1016/s0197-4580(02)00138-0

Galea, L.A.M., Roes, M.M., Dimech, C.J., Chow, C., Mahmoud, R., Lieblich, S.E., Duarte-Guterman, P., 2018. Premarin has opposing effects on spatial learning, neural activation, and serum cytokine levels in middle-aged female rats depending on reproductive history. Neurobiology of Aging 70, 291–307. https://doi.org/10.1016/j.neurobiolaging.2018.06.030

Gatewood, J.D., Morgan, M.D., Eaton, M., McNamara, I.M., Stevens, L.F., Macbeth, A.H., Meyer, E.A.A., Lomas, L.M., Kozub, F.J., Lambert, K.G., Kinsley, C.H., 2005. Motherhood mitigates aging-related decrements in learning and memory and positively affects brain aging in the rat. Brain Res Bull 66, 91–98. https://doi.org/10.1016/j.brainresbull.2005.03.016

Glynn, L.M., 2012. Increasing parity is associated with cumulative effects on memory. J Womens Health (Larchmt) 21, 1038–1045. https://doi.org/10.1089/jwh.2011.3206

Guillemin, G.J., Smith, D.G., Smythe, G.A., Armati, P.J., Brew, G.J., 2003. Expression of The Kynurenine Pathway Enzymes in Human Microglia and Macrophages, in: Allegri, G., Costa, C.V.L., Ragazzi, E., Steinhart, H., Varesio, L. (Eds.), Developments in Tryptophan and Serotonin Metabolism, Advances in Experimental Medicine and Biology. Springer US, Boston, MA, pp. 105–112. https://doi.org/10.1007/978-1-4615-0135-0_12

Gundersen, H.J.G., 1988. The nucleator. Journal of Microscopy 151, 3–21. https://doi.org/10.1111/j.1365-2818.1988.tb04609.x

Haim, A., Julian, D., Albin-Brooks, C., Brothers, H.M., Lenz, K.M., Leuner, B., 2017. A survey of neuroimmune changes in pregnant and postpartum female rats. Brain, Behavior, and Immunity 59, 67–78. https://doi.org/10.1016/j.bbi.2016.09.026

Hasegawa, T., Mikoda, N., Kitazawa, M., LaFerla, F.M., 2010. Treatment of Alzheimer’s disease with anti-homocysteic acid antibody in 3xTg-AD male mice. PLoS One 5, e8593. https://doi.org/10.1371/journal.pone.0008593

Hers, I., Vincent, E.E., Tavaré, J.M., 2011. Akt signalling in health and disease. Cell Signal 23, 1515–1527. https://doi.org/10.1016/j.cellsig.2011.05.004

Hoekzema, E., Barba-Müller, E., Pozzobon, C., Picado, M., Lucco, F., García-García, D., Soliva, J.C., Tobeña, A., Desco, M., Crone, E.A., Ballesteros, A., Carmona, S., Vilarroya, O., 2017. Pregnancy leads to long-lasting changes in human brain structure. Nat Neurosci 20, 287–296. https://doi.org/10.1038/nn.4458

Hong, S., Dissing-Olesen, L., Stevens, B., 2016. New insights on the role of microglia in synaptic pruning in health and disease. Curr Opin Neurobiol 36, 128–134. https://doi.org/10.1016/j.conb.2015.12.004

Hu, M., Becker, J.B., 2003. Effects of Sex and Estrogen on Behavioral Sensitization to Cocaine in Rats. J. Neurosci. 23, 693–699. https://doi.org/10.1523/JNEUROSCI.23-02-00693.2003

Hunt, C., Schenker, L., Kennedy, M., 1996. PSD-95 Is Associated with the Postsynaptic Density and Not with the Presynaptic Membrane at Forebrain Synapses. The Journal of neuroscience : the official journal of the Society for Neuroscience 16, 1380–8. https://doi.org/10.1523/JNEUROSCI.16-04-01380.1996

Huo, Y., Cheng, L., Wang, C., Deng, Y., Hu, R., Shi, L., Wan, Q., Chen, Lulu, Zeng, T., Yu, X., Tang, X., Yan, L., Qin, G., Chen, G., Gao, Z., Wang, G., Shen, F., Luo, Z., Qin, Y., Chen, Li, Li, Q., Ye, Z., Zhang, Y., Bi, Y., Lu, J., Li, M., Xu, M., Xu, Y., Wang, T., Zhao, Z., Chen, Y., Qi, H., Zhu, Y., Hu, C., Su, Q., Liu, C., Wang, Y., Wu, S., Yang, T., Deng, H., Zhao, J., Mu, Y., Ning, G., Wang, W., Lin, A., REACTION Group, 2021. Associations between parity, pregnancy loss, and breastfeeding duration and risk of maternal type 2 diabetes: An observational cohort study. J Diabetes 13, 857–867. https://doi.org/10.1111/1753-0407.13176

Jung, H.Y., Kim, W., Hahn, K.R., Kwon, H.J., Nam, S.M., Chung, J.Y., Yoon, Y.S., Kim, D.W., Yoo, D.Y., Hwang, I.K., 2020. Effects of Pyridoxine Deficiency on Hippocampal Function and Its Possible Association with V-Type Proton ATPase Subunit B2 and Heat Shock Cognate Protein 70. Cells 9, E1067. https://doi.org/10.3390/cells9051067

Kelleher, R.J., Govindarajan, A., Jung, H.-Y., Kang, H., Tonegawa, S., 2004. Translational Control by MAPK Signaling in Long-Term Synaptic Plasticity and Memory. Cell 116, 467–479. https://doi.org/10.1016/S0092-8674(04)00115-1

Kempermann, G., Gage, F.H., Aigner, L., Song, H., Curtis, M.A., Thuret, S., Kuhn, H.G., Jessberger, S., Frankland, P.W., Cameron, H.A., Gould, E., Hen, R., Abrous, D.N., Toni, N., Schinder, A.F., Zhao, X., Lucassen, P.J., Frisén, J., 2018. Human Adult Neurogenesis: Evidence and Remaining Questions. Cell Stem Cell 23, 25–30. https://doi.org/10.1016/j.stem.2018.04.004

Kim, E., Sheng, M., 2004. PDZ domain proteins of synapses. Nat Rev Neurosci 5, 771–781. https://doi.org/10.1038/nrn1517

Kim, P., Dufford, A.J., Tribble, R.C., 2018. Cortical thickness variation of the maternal brain in the first 6 months postpartum: associations with parental self-efficacy. Brain Struct Funct 223, 3267– 3277. https://doi.org/10.1007/s00429-018-1688-z

Kim, P., Leckman, J.F., Mayes, L.C., Feldman, R., Wang, X., Swain, J.E., 2010. The plasticity of human maternal brain: longitudinal changes in brain anatomy during the early postpartum period. Behav Neurosci 124, 695–700. https://doi.org/10.1037/a0020884

Kinsley, C.H., Trainer, R., Stafisso-Sandoz, G., Quadros, P., Marcus, L.K., Hearon, C., Meyer, E.A.A., Hester, N., Morgan, M., Kozub, F.J., 2006. Motherhood and the hormones of pregnancy modify concentrations of hippocampal neuronal dendritic spines. Hormones and Behavior 49, 131–142. https://doi.org/10.1016/j.yhbeh.2005.05.017

Klann, E., Dever, T.E., 2004. Biochemical mechanisms for translational regulation in synaptic plasticity. Nat Rev Neurosci 5, 931–942. https://doi.org/10.1038/nrn1557

Lange, A.G., Barth, C., Kaufmann, T., Anatürk, M., Suri, S., Ebmeier, K.P., Westlye, L.T., 2020. The maternal brain: Region-specific patterns of brain aging are traceable decades after childbirth. Hum Brain Mapp 41, 4718–4729. https://doi.org/10.1002/hbm.25152

Lecanu, L., Jay Waldron, Althea McCourty, 2010. Aging differentially affects male and female neural stem cell neurogenic properties. SCCAA 119. https://doi.org/10.2147/SCCAA.S13035

Lee, B.H., Richard, J.E., de Leon, R.G., Yagi, S., Galea, L.A.M., 2022. Sex Differences in Cognition Across Aging. Curr Top Behav Neurosci. https://doi.org/10.1007/7854_2022_309

Lemaire, V., Billard, J.-M., Dutar, P., George, O., Piazza, P.V., Epelbaum, J., Le Moal, M., Mayo, W., 2006. Motherhood-induced memory improvement persists across lifespan in rats but is abolished by a gestational stress. European Journal of Neuroscience 23, 3368–3374. https://doi.org/10.1111/j.1460-9568.2006.04870.x

Leuner, B., Glasper, E.R., Gould, E., 2010. Parenting and plasticity. Trends in Neurosciences 33, 465–473. https://doi.org/10.1016/j.tins.2010.07.003

Levitz, M., Young, B.K., 1977. Estrogens in pregnancy. Vitam Horm 35, 109–147. https://doi.org/10.1016/s0083-6729(08)60522-1

Lisofsky, N., Gallinat, J., Lindenberger, U., Kühn, S., 2019. Postpartal Neural Plasticity of the Maternal Brain: Early Renormalization of Pregnancy-Related Decreases? Neurosignals 27, 12–24. https://doi.org/10.33594/000000105

Luders, E., Gingnell, M., Poromaa, I.S., Engman, J., Kurth, F., Gaser, C., 2018. Potential Brain Age Reversal after Pregnancy: Younger Brains at 4–6 Weeks Postpartum. Neuroscience 386, 309–314. https://doi.org/10.1016/j.neuroscience.2018.07.006

Luders, E., Kurth, F., Gingnell, M., Engman, J., Yong, E.-L., Poromaa, I.S., Gaser, C., 2020. From baby brain to mommy brain: Widespread gray matter gain after giving birth. Cortex 126, 334–342. https://doi.org/10.1016/j.cortex.2019.12.029

Luppi, P., 2003. How immune mechanisms are affected by pregnancy. Vaccine 21, 3352–3357. https://doi.org/10.1016/S0264-410X(03)00331-1

Lynch, M.A., 2022. Exploring Sex-Related Differences in Microglia May Be a Game-Changer in Precision Medicine. Front. Aging Neurosci. 14, 868448. https://doi.org/10.3389/fnagi.2022.868448

Macbeth, A.H., Scharfman, H.E., MacLusky, N.J., Gautreaux, C., Luine, V.N., 2008. Effects of multiparity on recognition memory, monoaminergic neurotransmitters, and brain-derived neurotrophic factor (BDNF). Hormones and Behavior 54, 7–17. https://doi.org/10.1016/j.yhbeh.2007.08.011

Maes, M., Leonard, B.E., Myint, A.M., Kubera, M., Verkerk, R., 2011. The new ‘5-HT’ hypothesis of depression: Cell-mediated immune activation induces indoleamine 2,3-dioxygenase, which leads to lower plasma tryptophan and an increased synthesis of detrimental tryptophan catabolites (TRYCATs), both of which contribute to the onset of depression. Progress in Neuro-Psychopharmacology and Biological Psychiatry 35, 702–721. https://doi.org/10.1016/j.pnpbp.2010.12.017

Martínez-García, M., Paternina-Die, M., Barba-Müller, E., Martín de Blas, D., Beumala, L., Cortizo, R., Pozzobon, C., Marcos-Vidal, L., Fernández-Pena, A., Picado, M., Belmonte-Padilla, E., Massó- Rodriguez, A., Ballesteros, A., Desco, M., Vilarroya, Ó., Hoekzema, E., Carmona, S., 2021a. Do Pregnancy-Induced Brain Changes Reverse? The Brain of a Mother Six Years after Parturition. Brain Sci 11, 168. https://doi.org/10.3390/brainsci11020168

Martínez-García, M., Paternina-Die, M., Desco, M., Vilarroya, O., Carmona, S., 2021b. Characterizing the Brain Structural Adaptations Across the Motherhood Transition. Front Glob Womens Health 2, 742775. https://doi.org/10.3389/fgwh.2021.742775

McCormack, C., Callaghan, B.L., Pawluski, J.L., 2023. It’s Time to Rebrand “Mommy Brain.” JAMA Neurol. https://doi.org/10.1001/jamaneurol.2022.5180

Midttun, Ø., Hustad, S., Solheim, E., Schneede, J., Ueland, P.M., 2005. Multianalyte Quantification of Vitamin B6 and B2 Species in the Nanomolar Range in Human Plasma by Liquid Chromatography–Tandem Mass Spectrometry. Clinical Chemistry 51, 1206–1216. https://doi.org/10.1373/clinchem.2005.051169

Mor, G., Cardenas, I., 2010. The Immune System in Pregnancy: A Unique Complexity: IMMUNE SYSTEM IN PREGNANCY. American Journal of Reproductive Immunology 63, 425–433. https://doi.org/10.1111/j.1600-0897.2010.00836.x

Morris, M.S., Picciano, M.F., Jacques, P.F., Selhub, J., 2008. Plasma pyridoxal 5’-phosphate in the US population: the National Health and Nutrition Examination Survey, 2003-2004. Am J Clin Nutr 87, 1446–1454. https://doi.org/10.1093/ajcn/87.5.1446

Mouton, P.R., Long, J.M., Lei, D.-L., Howard, V., Jucker, M., Calhoun, M.E., Ingram, D.K., 2002. Age and gender effects on microglia and astrocyte numbers in brains of mice. Brain Research 956, 30–35. https://doi.org/10.1016/S0006-8993(02)03475-3

Napso, T., Yong, H.E.J., Lopez-Tello, J., Sferruzzi-Perri, A.N., 2018. The Role of Placental Hormones in Mediating Maternal Adaptations to Support Pregnancy and Lactation. Front Physiol 9, 1091. https://doi.org/10.3389/fphys.2018.01091

Ning, K., Zhao, L., Franklin, M., Matloff, W., Batta, I., Arzouni, N., Sun, F., Toga, A.W., 2020. Parity is associated with cognitive function and brain age in both females and males. Sci Rep 10, 6100. https://doi.org/10.1038/s41598-020-63014-7

Oatridge, A., Holdcroft, A., Saeed, N., Hajnal, J.V., Puri, B.K., Fusi, L., Bydder, G.M., 2002. Change in brain size during and after pregnancy: study in healthy women and women with preeclampsia. AJNR Am J Neuroradiol 23, 19–26.

Orchard, E.R., Rutherford, H.J.V., Holmes, A.J., Jamadar, S.D., 2023. Matrescence: lifetime impact of motherhood on cognition and the brain. Trends in Cognitive Sciences S1364661322003023. https://doi.org/10.1016/j.tics.2022.12.002

Orchard, E.R., Ward, P.G.D., Sforazzini, F., Storey, E., Egan, G.F., Jamadar, S.D., 2020. Relationship between parenthood and cortical thickness in late adulthood. PLoS One 15, e0236031. https://doi.org/10.1371/journal.pone.0236031

Parikh, N.I., Cnattingius, S., Dickman, P.W., Mittleman, M.A., Ludvigsson, J.F., Ingelsson, E., 2010. Parity and risk of later-life maternal cardiovascular disease. Am Heart J 159, 215–221.e6. https://doi.org/10.1016/j.ahj.2009.11.017

Pascual, Z.N., Langaker, M.D., 2022. Physiology, Pregnancy, in: StatPearls. StatPearls Publishing, Treasure Island (FL).

Pascual, Z.N., Langaker, M.D., 2021. Physiology, Pregnancy, StatPearls [Internet]. StatPearls Publishing.

Pawluski, J.L., Barakauskas, V.E., Galea, L. a. M., 2010. Pregnancy decreases oestrogen receptor alpha expression and pyknosis, but not cell proliferation or survival, in the hippocampus. J Neuroendocrinol 22, 248–257. https://doi.org/10.1111/j.1365-2826.2010.01960.x

Pawluski, J.L., Galea, L.A.M., 2007. Reproductive experience alters hippocampal neurogenesis during the postpartum period in the dam. Neuroscience 149, 53–67. https://doi.org/10.1016/j.neuroscience.2007.07.031

Pawluski, J.L., Walker, S.K., Galea, L.A.M., 2006. Reproductive experience differentially affects spatial reference and working memory performance in the mother. Hormones and Behavior 49, 143– 149. https://doi.org/10.1016/j.yhbeh.2005.05.016

Pertovaara, M., Raitala, A., Lehtimäki, T., Karhunen, P.J., Oja, S.S., Jylhä, M., Hervonen, A., Hurme, M., 2006. Indoleamine 2,3-dioxygenase activity in nonagenarians is markedly increased and predicts mortality. Mechanisms of Ageing and Development 127, 497–499. https://doi.org/10.1016/j.mad.2006.01.020

Plantone, D., Pardini, M., Rinaldi, G., 2021. Riboflavin in Neurological Diseases: A Narrative Review. Clin Drug Investig 41, 513–527. https://doi.org/10.1007/s40261-021-01038-1

Posillico, C.K., Schwarz, J.M., 2016. An investigation into the effects of antenatal stressors on the postpartum neuroimmune profile and depressive-like behaviors. Behavioural Brain Research 298, 218–228. https://doi.org/10.1016/j.bbr.2015.11.011

Qiu, W., Go, K.A., Lamers, Y., Galea, L.A.M., 2021. Postpartum corticosterone and fluoxetine shift the tryptophan-kynurenine pathway in dams. Psychoneuroendocrinology 130, 105273. https://doi.org/10.1016/j.psyneuen.2021.105273

Rehbein, E., Hornung, J., Sundström Poromaa, I., Derntl, B., 2021. Shaping of the Female Human Brain by Sex Hormones: A Review. Neuroendocrinology 111, 183–206. https://doi.org/10.1159/000507083

Reinken, L., Dapunt, O., 1978. Vitamin B6 nutriture during pregnancy. Int J Vitam Nutr Res 48, 341–347.

Robinson, D.P., Klein, S.L., 2012. Pregnancy and pregnancy-associated hormones alter immune responses and disease pathogenesis. Hormones and Behavior, Special Issue: The Neuroendocrine-Immune Axis in Health and Disease 62, 263–271. https://doi.org/10.1016/j.yhbeh.2012.02.023

Rossetti, M.F., Varayoud, J., Lazzarino, G.P., Luque, E.H., Ramos, J.G., 2016. Pregnancy and lactation differentially modify the transcriptional regulation of steroidogenic enzymes through DNA methylation mechanisms in the hippocampus of aged rats. Molecular and Cellular Endocrinology 429, 73–83. https://doi.org/10.1016/j.mce.2016.03.037

Sato, K., 2015. Effects of Microglia on Neurogenesis. Glia 63, 1394–1405. https://doi.org/10.1002/glia.22858

Schumacher, A., Costa, S.-D., Zenclussen, A.C., 2014. Endocrine Factors Modulating Immune Responses in Pregnancy. Front. Immunol. 5. https://doi.org/10.3389/fimmu.2014.00196

Sherer, M.L., Posillico, C.K., Schwarz, J.M., 2018. The psychoneuroimmunology of pregnancy. Front Neuroendocrinol 51, 25–35. https://doi.org/10.1016/j.yfrne.2017.10.006

Sherer, M.L., Posillico, C.K., Schwarz, J.M., 2017. An examination of changes in maternal neuroimmune function during pregnancy and the postpartum period. Brain Behav Immun 66, 201–209. https://doi.org/10.1016/j.bbi.2017.06.016

Speisman, R.B., Kumar, A., Rani, A., Pastoriza, J.M., Severance, J.E., Foster, T.C., Ormerod, B.K., 2013. Environmental enrichment restores neurogenesis and rapid acquisition in aged rats. Neurobiology of Aging 34, 263–274. https://doi.org/10.1016/j.neurobiolaging.2012.05.023

Tsokas, P., Grace, E.A., Chan, P., Ma, T., Sealfon, S.C., Iyengar, R., Landau, E.M., Blitzer, R.D., 2005. Local Protein Synthesis Mediates a Rapid Increase in Dendritic Elongation Factor 1A after Induction of Late Long-Term Potentiation. J. Neurosci. 25, 5833–5843. https://doi.org/10.1523/JNEUROSCI.0599-05.2005

Tsokas, P., Ma, T., Iyengar, R., Landau, E.M., Blitzer, R.D., 2007. Mitogen-Activated Protein Kinase Upregulates the Dendritic Translation Machinery in Long-Term Potentiation by Controlling the Mammalian Target of Rapamycin Pathway. J. Neurosci. 27, 5885–5894. https://doi.org/10.1523/JNEUROSCI.4548-06.2007

van de Kamp, J.L., Smolen, A., 1995. Response of kynurenine pathway enzymes to pregnancy and dietary level of vitamin B-6. Pharmacology Biochemistry and Behavior 51, 753–758. https://doi.org/10.1016/0091-3057(95)00026-S

Veen, C., Myint, A.M., Burgerhout, K.M., Schwarz, M.J., Schütze, G., Kushner, S.A., Hoogendijk, W.J., Drexhage, H.A., Bergink, V., 2016. Tryptophan pathway alterations in the postpartum period and in acute postpartum psychosis and depression. Journal of Affective Disorders 189, 298–305. https://doi.org/10.1016/j.jad.2015.09.064

Wang, C., Yue, H., Hu, Z., Shen, Y., Ma, J., Li, J., Wang, X.-D., Wang, Liang, Sun, B., Shi, P., Wang, Lang, Gu, Y., 2020. Microglia mediate forgetting via complement-dependent synaptic elimination. Science 367, 688–694. https://doi.org/10.1126/science.aaz2288

Wichers, M.C., Koek, G.H., Robaeys, G., Verkerk, R., Scharpé, S., Maes, M., 2005. IDO and interferon-α-induced depressive symptoms: a shift in hypothesis from tryptophan depletion to neurotoxicity. Mol Psychiatry 10, 538–544. https://doi.org/10.1038/sj.mp.4001600

Workman, J.L., Gobinath, A.R., Kitay, N.F., Chow, C., Brummelte, S., Galea, L.A.M., 2016. Parity modifies the effects of fluoxetine and corticosterone on behavior, stress reactivity, and hippocampal neurogenesis. Neuropharmacology 105, 443–453. https://doi.org/10.1016/j.neuropharm.2015.11.027

Yoshii, A., Constantine-Paton, M., 2014. Postsynaptic localization of PSD-95 is regulated by all three pathways downstream of TrkB signaling. Frontiers in Synaptic Neuroscience 6.

Zhou, Y., Su, Y., Li, S., Kennedy, B.C., Zhang, D.Y., Bond, A.M., Sun, Y., Jacob, F., Lu, L., Hu, P., Viaene, A.N., Helbig, I., Kessler, S.K., Lucas, T., Salinas, R.D., Gu, X., Chen, H.I., Wu, H., Kleinman, J.E., Hyde, T.M., Nauen, D.W., Weinberger, D.R., Ming, G., Song, H., 2022. Molecular landscapes of human hippocampal immature neurons across lifespan. Nature 607, 527–533. https://doi.org/10.1038/s41586-022-04912-w

Zysset-Burri, D.C., Bellac, C.L., Leib, S.L., Wittwer, M., 2013. Vitamin B6 reduces hippocampal apoptosis in experimental pneumococcal meningitis. BMC Infectious Diseases 13, 393. https://doi.org/10.1186/1471-2334-13-393

